# Rampant transcription replication conflict creates therapeutic vulnerability in extrachromosomal DNA containing cancers

**DOI:** 10.1101/2024.03.29.586681

**Authors:** Jun Tang, Natasha E. Weiser, Guiping Wang, Sudhir Chowdhry, Ellis J. Curtis, Yanding Zhao, Ivy Tsz-Lo Wong, Georgi K. Marinov, Rui Li, Philip Hanoian, Edison Tse, Ryan Hansen, Joshua Plum, Auzon Steffy, Snezana Milutinovic, S. Todd Meyer, Christina Curtis, William J. Greenleaf, Vineet Bafna, Stephen J. Benkovic, Anthony B. Pinkerton, Shailaja Kasibhatla, Christian A. Hassig, Paul S. Mischel, Howard Y. Chang

**Affiliations:** Department of Pathology, Stanford University School of Medicine, Stanford, CA, USA; Sarafan ChEM-H, Stanford University, Stanford, CA, USA; Center for Personal Dynamic Regulomes, Stanford University, Stanford, CA, USA; Boundless Bio, San Diego, CA, USA; Department of Genetics, Stanford University School of Medicine, Stanford, CA, USA; Department of Chemistry, Pennsylvania State University, University Park, PA, USA; Department of Computer Science and Engineering, University of California, San Diego, La Jolla, CA, USA; Medical Scientist Training Program, University of California, San Diego, La Jolla, CA, USA; Department of Dermatology, Stanford University School of Medicine, Stanford, CA, USA; Department of Medicine, Stanford University School of Medicine, Stanford, CA, USA; Stanford Cancer Institute, Stanford University School of Medicine, Stanford, CA, USA; Howard Hughes Medical Institute, Stanford University School of Medicine, Stanford, CA, USA

## Abstract

Extrachromosomal DNA (ecDNA) presents a major challenge for precision medicine, contributing to poor survival for patients with oncogene-amplified tumours. EcDNA renders tumours resistant to targeted treatments by facilitating massive transcription of oncogenes and rapid genome evolution. At present, there are no ecDNA- specific treatments. Here we show that enhancing transcription replication conflict enables targeted elimination of ecDNA-containing cancers, exposing an actionable vulnerability. Stepwise analyses of ecDNA transcription reveal landscapes of pervasive RNA transcription and associated single-stranded DNA, leading to excessive transcription replication conflicts and replication stress (RS) compared to chromosomal loci. Nucleotide incorporation onto growing DNA strands is markedly slower on ecDNA, and RS is significantly higher in ecDNA-containing tumours regardless of cancer type or oncogene cargo. Replication Protein A2 phosphorylated on serine 33, a mediator of DNA damage repair that binds single-stranded DNA, shows elevated localization on ecDNA in a transcription dependent manner, along with increased DNA double strand breaks, and activation of the S-phase checkpoint kinase, CHK1. Genetic or pharmacological CHK1 inhibition abrogates the DNA replication check point, causing extensive and preferential tumour cell death in ecDNA-containing tumours as they enter S-phase. To exploit this vulnerability, we develop a highly selective, potent, and bioavailable oral CHK1 inhibitor, BBI-2779, and demonstrate that it preferentially kills ecDNA-containing tumour cells. In a gastric cancer model containing *FGFR2* on ecDNA, BBI-2779, suppresses tumour growth and prevents ecDNA-mediated acquired resistance to the pan-FGFR inhibitor infigratinib, resulting in potent and sustained tumour regression in mice. These results reveal transcription-replication conflict as an ecDNA-generated vulnerability that can be targeted as an ecDNA-directed therapy and suggest that synthetic lethality of excess can be exploited as a strategy for treating cancer.

---

Extrachromosomal DNAs (ecDNAs) are a frequent mechanism for oncogene amplification in diverse cancer types and are associated with worse patient outcomes than other kinds of focal amplification^1,2^. EcDNAs can arise during the transition to, development, and progression of cancers and they exhibit unique biological features that provide fitness advantages to malignant cells^3^. The acentric structure of ecDNA facilitates random segregation, highly elevated copy number, intratumoural genetic heterogeneity, and rapid tumour evolution^1,4,5^, contributing to aggressive tumour growth and therapeutic resistance^6,7^. The circular topology of ecDNAs also profoundly alters transcription^8,9^. EcDNAs exhibit highly accessible chromatin and increased oncogene expression compared to non-circular amplifications, even after controlling for DNA copy number^1,10–12^. Further, ecDNAs can cluster in the nucleus to generate new, functional enhancer-promoter interactions both in *cis* and in *trans*^10,12^. Prior studies showed that ecDNA highly transcribe annotated protein-coding genes^11^, but it is unclear whether the full landscape of RNA transcription--such as intergenic, antisense, or other long noncoding RNAs--is altered. EcDNA exhibits open chromatin and is marked by active histone modifications such as H3K27ac and H3K4me3^1,11,13,14^, raising the possibility of a more permissive transcriptional environment. We hypothesized that the highly accessible chromatin of ecDNA could generate a therapeutically exploitable vulnerability.

## Landscape of ecDNA transcription reveals pervasive RNA production and associated ssDNA

To test this hypothesis, we performed Global Run-On sequencing (GRO-seq^16^) and rRNA-depleted RNA sequencing (Ribo-Zero) to profile nascent transcription and accumulated RNAs respectively (**Fig. 1a**), providing a comprehensive landscape of RNA biogenesis from ecDNAs. To control for the effects of focal amplification and assess ecDNA-specific transcriptional changes, we focused on a pair of isogenic colorectal cancer cell lines derived from the same patient: COLO320DM (*MYC* amplification on ecDNA) and COLO320HSR (chromosomal *MYC* amplification or homogeneously staining region; HSR) that are nearly matched for amplicon copy number as revealed by whole genome sequencing (WGS)^11^. Notably, COLO320DM showed approximately four-fold increase in nascent RNA and accumulated RNA read density from ecDNA, beyond the level expected from differences in amplicon copy number compared to COLO320HSR (**Fig. 1b**).

**Figure 1:**
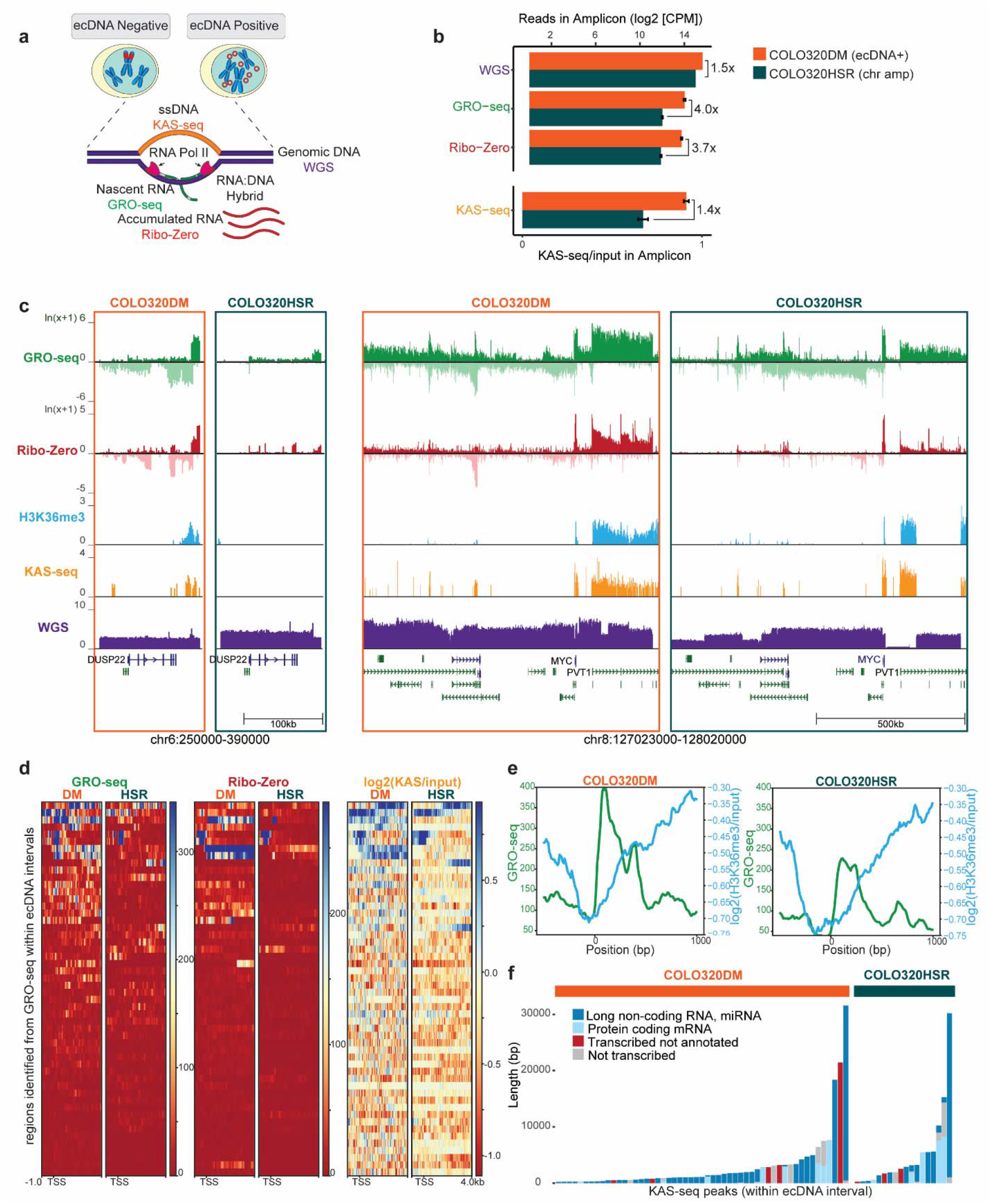
Pervasive transcription on ecDNA drives ssDNA accumulation. (**a**) Schematic of relevant genomic assays. (**b**) Read density of genomic assays in COLO320DM and COLO320HSR within the ecDNA interval in total counts per million (CPM). KAS-seq read density is shown as (CPM) of the KAS-seq relative to CPM of the input. For GRO-seq, Ribo- Zero, and KAS-seq the mean of two biological replicates is shown, error bars indicate standard deviation. A single replicate is shown for WGS (**c**) Genome tracks highlighting 2 regions within the ecDNA interval. H3K36me3 ChIP-seq and KAS-seq signal are displayed as log2 of input- normalized coverage. One representative biological replicate for each condition is visualized. (**d**) Metagene heatmap plot visualization of GRO-seq, Ribo Zero RNA-seq, and log2 of input- normalized coverage of KAS-seq within the ecDNA interval. All plots are anchored at the GRO- seq TSS as identified by HOMER. One representative biological replicate for each condition is visualized. (**e**) Metagene plot showing GRO-seq and H3K36me3 ChIP-seq coverage within the ecDNA interval. One representative biological replicate for each condition is visualized. All plots are anchored at the GRO-seq TSS as identified by HOMER using both biological replicates. H3K36me3 ChIP-seq coverage is displayed as log2(H3K36me3 IP/input). (**f**) KAS- seq peaks from two biological replicates in the ecDNA interval annotated by transcription status according to GRO-seq data and annotation status according to Gencode v43.

The increase in transcription was not limited to the *MYC* oncogene but was pervasive across the entire ecDNA, including noncoding, antisense, and numerous previously unannotated transcripts (**Fig. 1c, d**). This widespread increase in transcription is specific to the ecDNA, as GRO-seq and Ribo-Zero read densities on chromosomes were comparable between COLO320DM and COLO320HSR (**Extended Fig. 1**). We performed *de novo* transcript identification within the amplicon intervals using GRO-seq data and compared the same regions in COLO320DM versus COLO320HSR. We observed increases of both nascent and accumulated transcripts in COLO320DM compared to COLO320HSR, confirming that the increased transcription from ecDNA is amplicon-wide and not driven by a small number of differentially expressed transcripts (**Fig. 1d**). EcDNA also show increased occupancy of H3K36me3, a histone mark associated with RNA polymerase II elongation, downstream of transcription start sites (TSS) identified in our *de novo* transcript calling, providing orthogonal validation of rampant transcription (**Fig. 1c, e, Extended Fig. 2a**).

**Figure 2.**
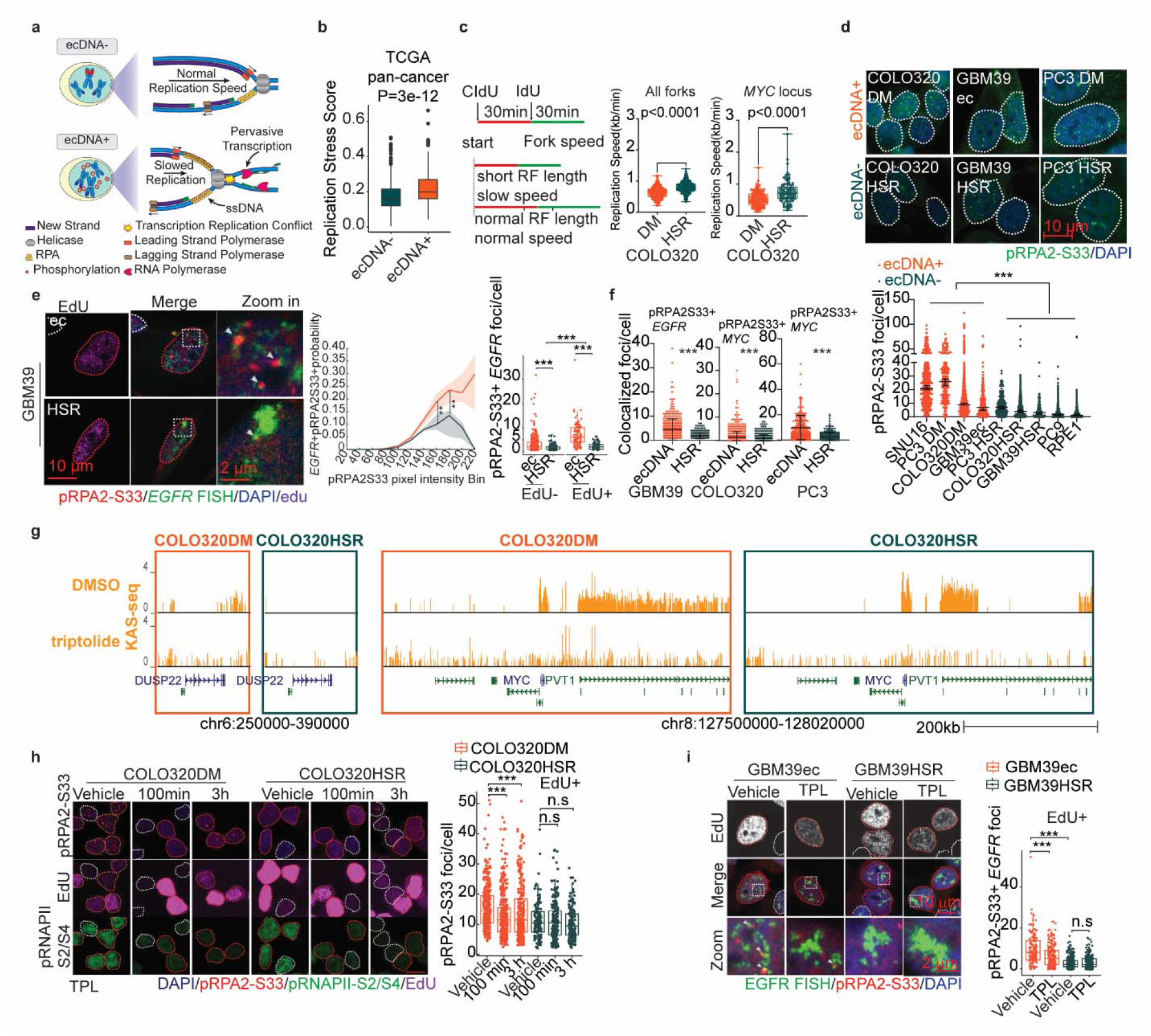
Transcription replication conflict creates replication stress on ecDNAs. (**a**) Schematics depict transcription replication conflict and replication stress. (**b**) Replication stress score computed in TCGA patients grouped by with or without ecDNA amplification (240 ecDNA positive, 639 ecDNA negative, which includes 110 linear, 529 no amplification, p values determined by two-sided Wilcoxon test. box center line, median; box limits, 25 and 75 quartiles; box whiskers, 1.5× interquartile range.) (**c**) DNA fiber assay combined with MYC FISH staining in COLO320DM and COLO320HSR cells. Left: Idu/Cldu labeling timeline and schematics depict DNA fiber length to infer replication fork progression rate. Middle: Global replication fork progression rate. Right: Replication fork progression rate at *MYC* locus. (box center line, median; box limits, 25th and 75th percentile; box whiskers, min to max, unpaired Kolmogorov- Smirnov test, All replication forks COL0320DM: n=348; COLO320HSR: n= 317; replication fork at *MYC* locus, COLO320DM: n= 143; COLO320HSR: n= 101). (**d**) Upper: Representative images of pRPA2-S33 immunofluorescence staining in 3 isogenic tumour cell line pairs. White dotted lines mark the boundary of cell nucleus as identified by DAPI staining; Lower, pRPA2-S33 foci number in individual cells, each dot indicated one cell. (Median with 95 CI, p values quantified by ordinary one-way ANOVA with multiple comparisons test, SNU16, n=666; PC3- DM, n= 205; COLO320DM, n=1191; GBM39ec, n=369; PC3-HSR, n=244; COLO320HSR, n=655; GBM39HSR, n=244; PC9, n=1505; RPE1, n=775.) (**e**) pRPA2-S33 immunofluorescence combined with *EGFR* FISH staining in GBM39ec and GBM39HSR cells, EdU was added 30 min before fixation. Left: Representative images. White dotted lines mark EdU- nucleus, while red dotted lines marks EdU+ nucleus; co-localized foci between pRPA2-S33 and *EGFR* were indicated by white arrow. Middle: proportion of pixels with pRPA2-S33 and EGFR colocalization within each pRPA2-S33 pixel intensity bins, data presented as median ± 25% quantile range (GBM39ec, n= 10; GBM39HSR, n=6). Right, *EGFR* foci number co-localized with pRPA2-S33; right, % of *EGFR* co-localized with pRPA2-S33. (Box plot parameters same as in Fig 1b, Two-tailed Mann-Whitney test, EdU- group: GBM39ec, n=299, GBM39HSR, n=201; EdU+ group: GBM39ec, n=120, GBM39HSR, n=83). (**f**) Quantification of the pRPA2- S33 colocalized with the amplified oncogenes in 3 isogenic cell line pairs: GBM39ec/GBM39HSR: EGFR, COLO320DM/COLO320HSR: MYC, PC3-DM/PC3-HSR: MYC. (mean ± SD, two tailed Mann-Whitney test, n number from left to right: 419, 284, 1085, 596, 209, 242). (**g**) Genome tracks highlighting 2 regions within the ecDNA interval in COLO320DM and COLO320HSR cells treated with 1 µM triptolide or vehicle for 3 hours. KAS-seq signal is displayed as log2 of input-normalized coverage. (**h**) COLO320DM and COLO320HSR cells were treated with 1 µM triptolide (TPL) for indicated time and EdU was added 30 min before fixing cells for pRPA2-S33 and pRNAPolII-S2/S4 immunofluorescence staining. Left, representative images in each group, red dotted line marks EdU+ nuclei and white dotted line marks EdU- nuclei; right, pRPA2-S33 foci number in EdU+ cells. (Box plot parameters same as in Fig 1b, two-sided Wilcoxon test, COLO320DM group: vehicle, n= 354, 100 min, n=350, 3 h, n= 269; COLO320HSR group: vehicle, n=130, 100 min, n= 185, 3 h, n= 161). (**i**) pRPA2-S33 immunofluorescence combined with EdU and *EGFR* FISH staining in GBM39ec and GBM39HSR cells treated with triptolide (20 µM) for 4hrs. Left, representative images in each group, red dotted line marks EdU+ nuclei and white dotted line marks EdU- nuclei, co-localized foci between pRPA2-S33 and *EGFR* were marked by white arrows. Right, quantification of *EGFR* foci number co-localized with pRPA2-S33 in replicating EdU+ cells. (Box plot parameters same as in Fig 1b, two-sided Wilcoxon test, GBM39ec vehicle, n=138, GBM39ec TPL, n=189, GBM39HSR vehicle, n=253, GBM39HSR TPL, n= 208) (* represents p<0.05, ** represents p<0.01, *** represents p<0.001, scale bar represents 10 µm or as otherwise specified in the image)

Elevated transcription is associated with single-stranded DNA (ssDNA) accumulation, due to the process of transcription itself, R loop formation from RNA:DNA hybrids, and transcription-replication conflict. To assess the influence of pervasive transcription on ecDNA structure, we performed kethoxal-assisted single-stranded DNA sequencing (KAS-seq)^17,18^ to map single-stranded DNA (ssDNA) genome-wide. After normalizing to input to account for copy number differences, we observe a 1.4-fold increase in KAS-seq read density within the ecDNA amplicon in COLO320DM compared to COLO320HSR **(Fig 1b, c**). The ssDNA regions on ecDNA extend from hundreds to over 20,000 basepairs (bp) and the majority of KAS-seq peaks overlap with transcribed regions, such as annotated noncoding transcripts (lncRNAs, miRNAs, 60%) and novel transcripts identified in GRO-seq (18%, **Fig. 1d, f, Extended Fig. 2b**). Taken together, these results suggest that ecDNAs provide a permissive chromatin environment for pervasive transcription initiation, leading to accumulated RNA species and ssDNA.

## Transcription replication conflict creates replication stress on ecDNAs

Pervasive transcription on ecDNA increases the possibility of transcription replication conflict. When RNA polymerase II collides with the DNA replication machinery, progression of the replication fork is stalled, incorporation of new nucleotides is slowed, ssDNA behind the replication fork is exposed and bound by phosphorylated RPA protein, and the cell experiences replication stress^19^ (**Fig. 2a**). This hypothesis predicts that ecDNA-containing cancer cells should have elevated DNA replication stress, and that the replication stress will be relieved by limiting transcription. First examining ecDNA-containing primary tumours, we grouped tumours from The Cancer Genome Atlas (TCGA) tumour patients into ecDNA-positive vs. negative cohorts based on WGS data analyzed by AmpliconArchitect^20^. We computed the replication stress score through a gene expression signature identified by Llobet.et al^21^ and found a significantly higher replication stress score in ecDNA-containing tumour patients (**Fig. 2b**). This result indicates that increased replication stress may be a common feature shared by ecDNA+ cancers. Next, conflicts between transcriptional and replicative machinery should lead to slower replication fork progression. To assay replication fork dynamics, we combined a DNA-fiber assay with DNA FISH to analyze replication fork dynamics in *MYC* amplified isogenic COLO320DM vs. COLO320HSR cells. The length of each Idu/CIdu track represents the velocity of each replication fork. We observed a slower replication fork progression rate in COLO320DM compared with COLO320HSR cells; importantly, double labeling of thymidine analog incorporation and *MYC* DNA FISH showed ecDNA had significantly slower replication fork progression compared to the same sequence on the chromosome (**Fig. 2c**).

To directly visualize replication stress in individual tumour cells and to determine its subnuclear localization, we detected RPA2 protein phosphorylation on serine 33 (pRPA2-S33) by immunofluorescence (IF). We analyzed pRPA2-S33 in a panel of cell lines including 3 isogenic cell line pairs: COLO320DM/COLO320HSR (*MYC*-amplified colorectal cancer), GBM39ec/GBM39HSR (*EGFR*-amplified glioblastoma), and PC3-DM/PC3-HSR (*MYC*- amplified prostate cancer), along with several other cell lines with or without ecDNA. Within each isogenic cell line pair, the amplified oncogene is shared but differs in its location on ecDNA or on a chromosome/HSR. We detected two- to three-fold higher pRPA2-S33 foci in ecDNA+ compared with ecDNA- tumour cells, indicating increased replication stress in ecDNA- containing tumour cells (**Fig. 2d**).

To determine whether replication stress is preferentially elevated on ecDNA and to determine whether it is enhanced on actively replicating DNA, we performed DNA FISH targeting the EGFR oncogene amplified on ecDNA, concurrently with EdU labeling, and IF detection of pRPA2-S33 in GBM39ec cells. We also examined these features in the isogenic counterpart, GBM39HSR in which amplified *EGFR* has reintegrated at the same copy number into chromosomes^11^. As hypothesized, we detected significantly higher replication stress on ecDNA, as measured by colocalization of pRPA2-S33 and *EGFR* FISH signal compared to GBM39HSR tumour cells, especially in EdU positive cells. Notably, in pixels with increasing pRPA2-S33 intensity, this colocalization ratio is more than tripled on ecDNA as opposed to HSR, which suggests specific molecular interactions rather than just spatial organization differences between oncogenes on ecDNA and HSR (**Fig. 2e**). We continued to observe increased pRPA2-S33 signal on *EGFR* in ecDNA+ cells after accounting for the total *EGFR* FISH signal, confirming that the higher replication stress on ecDNA compared to chromosomal amplification is not driven by differences in copy number (**Extended Fig. 3a**). pRPA2-S33 IF combined with DNA FISH staining in two other near-isogenic cell line pairs containing *MYC* amplifications, COLO320DM/COLO320HSR and PC3DM/PC3HSR, also showed higher replication stress on ecDNA compared with HSR (**Fig. 2f**). Our results across multiple cancer cell-types agnostic to the identity of the amplified oncogene collectively suggest that higher replication stress is a common feature of ecDNAs (**Extended Fig. 3b-d**).

**Figure 3.**
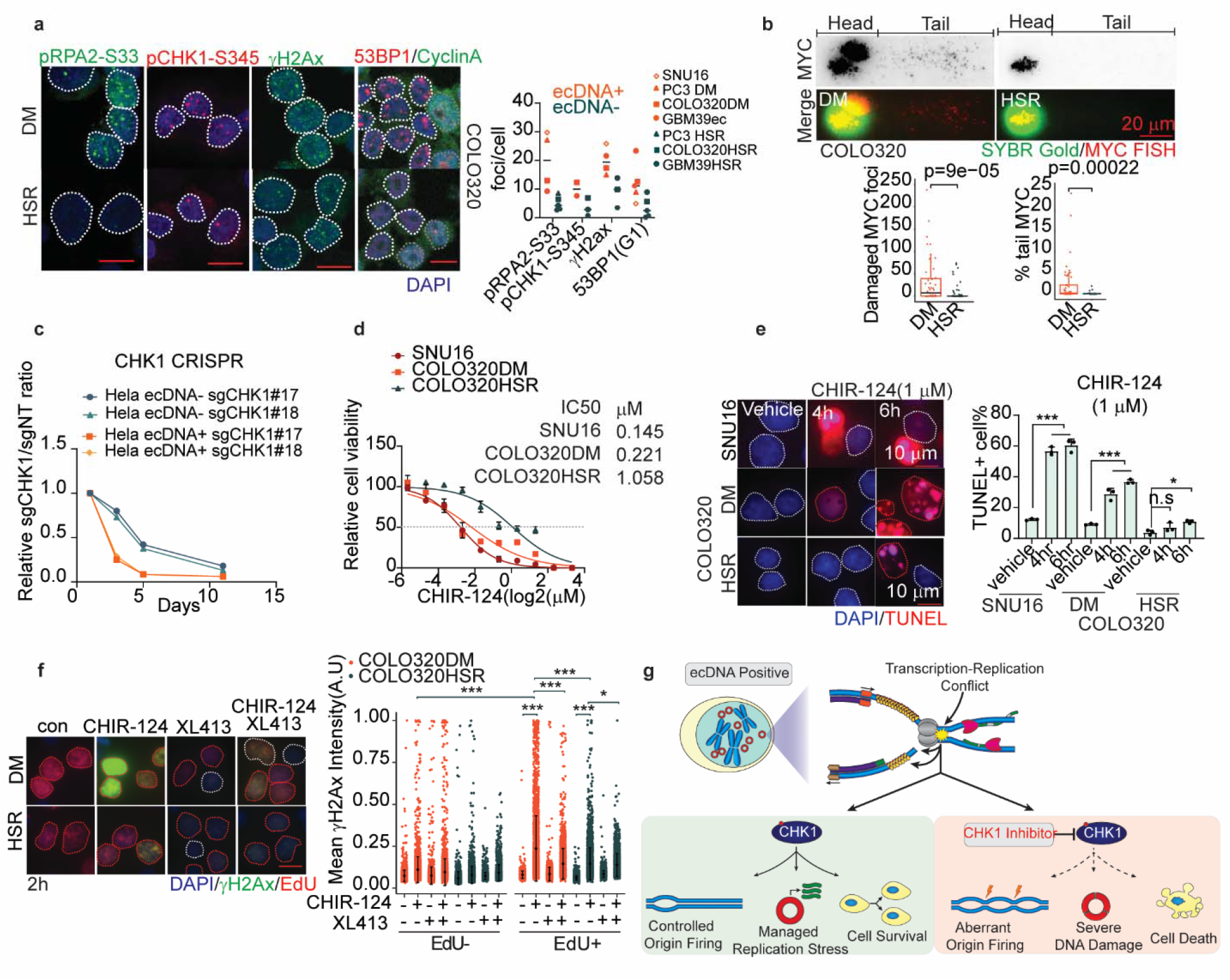
Replication stress activated S phase checkpoint and generated vulnerability to CHK1 inhibition in ecDNA-containing tumour cells. (**a**) Detection of pRPA2-S33, γH2AX, pCHK1-S345 and 53BP1/CyclinA in multiple cancer cell lines with or without amplicon on ecDNA. Left, representative images in COLO320DM and COLO320HSR cells. For pRPA2-S33, γH2AX and pCHK1-S345 IF staining, white dotted line mark the nucleus boundary; for 53BP1/CyclinA IF staining, white dotted line mark the G1 phase cell with Cyclin A negative staining and grey dotted line mark the Cylin A positive nucleus; Right, mean foci number in individual cell lines, 53BP1 foci number was quantified in G1 phase cells. Line indicates median, each dot indicates mean foci number in each cell line. (**b**) Comet- FISH assay in COLO320DM and COLO320HSR cells. Upper: representative images; lower: left, *MYC* foci number present in tail; right, percentage of *MYC* present in comet-tail normalized to whole cell. (Box plot parameters are the same as in Fig 1b, two-sided Wilcoxon test, *MYC* foci counting: COLO320DM, n=47, COLO320HSR, n= 60; % *MYC* tail: COLO320DM, n= 49, COLO320HSR, n= 33). (**c**) Relative cell number of Hela ecDNA+ and Hela ecDNA- cells transduced with sgRNAs targeting CHK1 to that with NT sgRNA for different time. (**d**) Cell viability curves of SNU16, COLO320DM and COLO320HSR in response to CHK1 inhibitor CHIR-124 for 3 days. (n=4, data presented as mean ± SD). (**e**) Terminal deoxynucleotidyl transferase dUTP nick end labeling (TUNEL) assay in cells subjected to 1 µM CHIR-124 for indicated time. Left, representative images, white dotted lines mark nuclei boundary; right, TUNEL positive cells percentage. (Data presented as mean ± SD, Ordinary one-way ANOVA with multiple comparison test, n=3). (**f**) γH2AX immunofluorescence staining in COLO320DM and COLO320HSR cells treated with CHIR-124 (100 nM) for 2 hours with or without the combination of CDC7i (XL413 20 µM), EdU was added 30min before fixation. Left, representative images, red dotted lines mark EdU+ nuclei and white dotted lines mark EdU- nuclei. Right, mean γH2AX intensity (arbitrary units) with different treatment. (mean ± SD, each dot indicates one cell, n number from left to right: EdU-: 3074, 4246, 3291, 4742, 3101, 3770, 2608, 2091; EdU+: 2428, 2859, 2909, 2890, 3346, 3491, 3232, 2060; two-tailed student’s t test). (**g**) Schematics depicting CHK1 activation in response to replication stress, which sensitizes ecDNA-containing tumour cells to targeted CHK1 inhibition through unscheduled replication origin firing and accumulation of excessive DNA damage, thus leads to cell death eventually. (* represents p<0.05, ** represents p<0.01, *** represents p<0.001, scale bar represents 10 µm or as otherwise specified in the image)

Having shown that ecDNAs have more open chromatin, increased transcription, and elevated replication stress, we set out to determine whether the elevated replication stress on ecDNA is a direct and potentially actionable consequence of pervasive transcription generated by ecDNA’s topology. We treated COLO320DM and COLO320HSR cells with triptolide, which inhibits transcription initiation through binding to the XPB subunit of the transcription factor complex TFIIH^22^. Active RNA polymerase II detected by IF showed that triptolide treatment significantly decreased transcriptional activity (**Extended Fig. 4a**). KAS-seq analysis in COLO320DM and COLO320HSR cells treated with triptolide revealed drastic reduction in ssDNA signals across the ecDNA amplicon, (**Fig. 2g**). We found that triptolide treatment significantly decreased pRPA2-S33 foci in COLO320DM cells, with negligible effect in COLO320HSR cells (**Fig. 2h**), suggesting that transcription contributes to the elevated replication stress in COLO320DM cells. In the GBM39 isogenic model, nascent transcription is modestly higher in GBM39ec than GBM39HSR as indicated by GRO-seq (**Extended Fig. 5a-b**). Triptolide treatment of GBM39ec cells significantly decreased replication stress on ecDNA as detected by combined pRPA2-S33 IF and *EGFR* FISH, whereas no obvious difference was observed on HSR (**Fig 2i, Extended Fig. 4b**). Taken together, our results demonstrate that ecDNAs exhibit higher levels of replication stress than chromosomal loci and that this increased replication stress is driven in large part by transcription.

**Figure 4.**
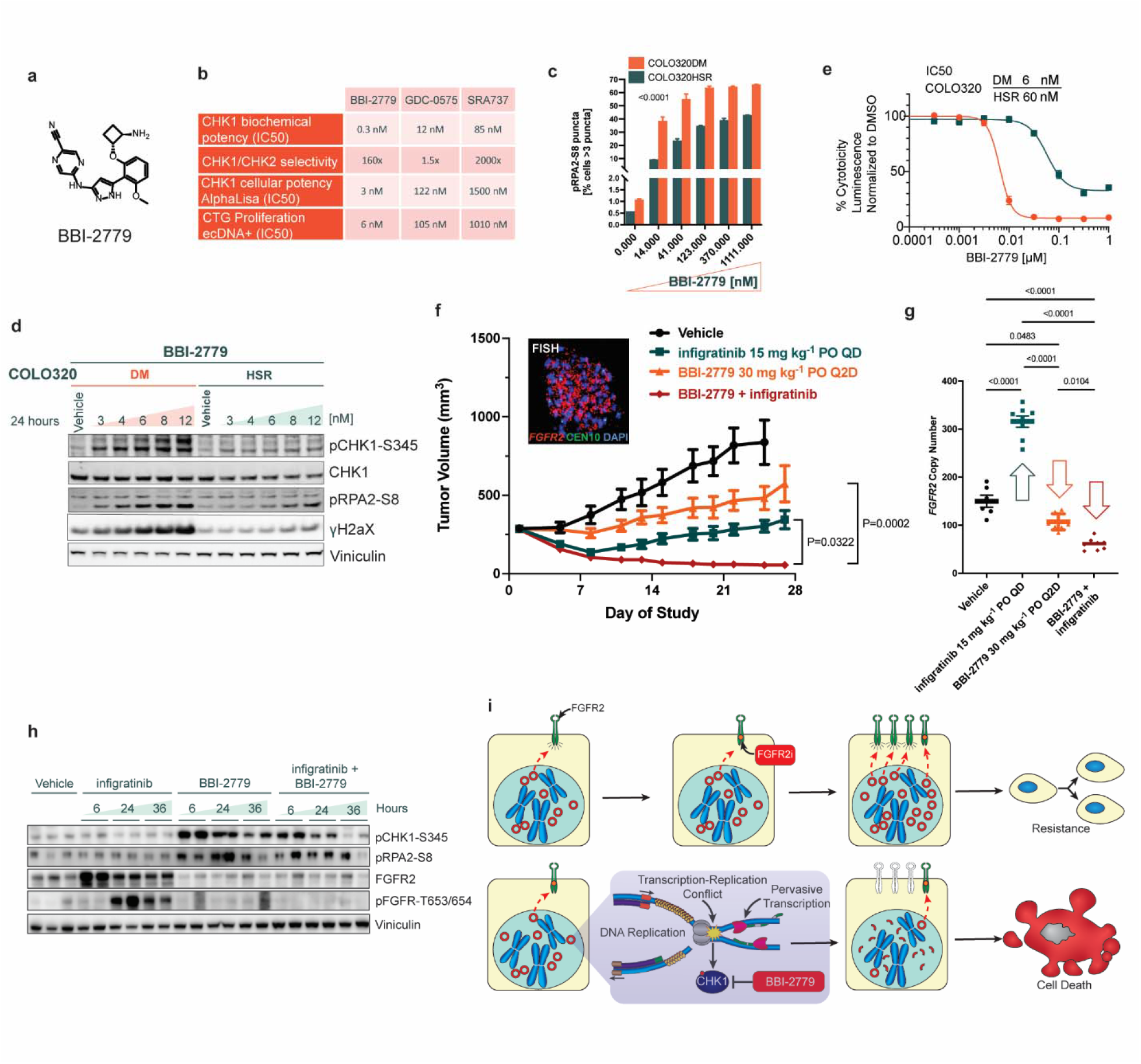
Oral CHK1i in combination with a pan-FGFRi demonstrates synergistic antitumour activity, and inhibits acquired resistance to targeted therapy manifested by ecDNA. (**a**) Chemical structure of BBI-2779. (**b**) In vitro and cellular potency of BBI-2779, and of reference compounds. AlphaLisa pCHK1-S345 activity was assessed in HT29 cells, while anti- proliferation potency was evaluated in COLO320DM cells. (**c-d**) Dose dependent induction of replication stress and that of the associated biomarkers upon treatment with BBI-2779 was investigated by measuring levels of hyperphosphorylated form of RPA32 Ser8, detected by immunofluorescence as well as by immunoblots in 4d. Significance was determined using ordinary 2-way ANOVA. (**e**) Differential tumour cell anti-proliferation activity of BBI-2779 was evaluated in COLO320DM and COLO320HSR tumour cells in a 5 day cell proliferation assay. (**f**) Embedded FISH image of SNU16 cells demonstrating *FGFR2*-positive ecDNA. SNU-16 cells were grown as tumour xenografts in SCID beige mice. After tumour establishment (∼285Lmm^3^), mice were treated with vehicle, BBI-2779 (30 mg kg−1), infigratinib (15 mg kg−1) or BBI-2779 (30 mg kg−1) plus infigratinib (15 mg kg−1), for 25 day (vehicle) or 27 days (other arms). Mean tumour volumes ± standard error of the mean are shown (n = 8 mice per group; one vehicle tumour was taken down on day 22 mouse was sacrificed due to large tumour volume). (**g**) FGFR2 copy number was evaluated by qPCR on tumour DNA. (**h**) Immunoblots of tumour lysates measuring elevated RS, DNA damage, and abrogation of oncoprotein FGFR2 expression. Significance was determined by one-way ANOVA with Tukey’s multiple comparisons. (**i**) ecDNA amplified oncogenes are hyper- transcribed, resulting in elevated RS, and reliance on CHK1 to manage DNA replication to maintain oncoprotein overexpression, and proliferation. Inhibition of CHK1 results in uncontrolled origin firing and failed cell cycle checkpoints, exacerbating RS in ecDNA-enabled tumour cells. Synthetic lethality to CHK1i in ecDNA+ oncogene amplified tumour cells is synergistic with targeted therapy resulting in enhanced cytotoxicity.

## Replication stress induces DNA damage on ecDNA

Replication stress contributes to endogenous DNA damage because stalled replication forks are unstable and prone to breakage, generating DNA lesions^23^. Therefore, we hypothesized that ecDNA-containing tumour cells may have higher baseline levels of DNA damage. In a panel of ecDNA+ and ecDNA- cancer cell lines, including three near-isogenic cell line pairs we found that in addition to having more pRPA2-S33 foci, ecDNA+ cells showed an average increased number of γH2AX and 53BP1 foci than the corresponding isogenic HSR and/or other ecDNA- cell lines (**Fig. 3a, Extended Fig. 6b-c**). Combined γH2AX IF with DNA FISH staining in isogenic cell line pairs confirmed enhanced DNA damage on ecDNAs, compared to chromosomal amplicons (**Extended Fig. 6d-e**). To further confirm the presence of DNA damage on ecDNA itself, we performed an alkaline-comet assay combined with *MYC* FISH staining in COLO320DM and COLO320HSR cells. In this assay, damaged DNA appears in the tail region of the comet. We observed significantly more *MYC* foci in the tail region of COLO320DM cells compared to COLO320HSR cells (**Fig. 3b**), which have comparable amplicon copy number. Taken together, these data demonstrate elevated DNA damage on ecDNA, relative to the same loci amplified on chromosomes. Thus, ecDNA-containing cancer cells may be hyper-reliant on the RS regulation machinery to cope with the elevated levels of baseline DNA damage driven by transcription replication conflicts.

## Vulnerability to CHK1 inhibitors in cancer cells with ecDNA

We reasoned that this unique feature of ecDNA, might generate an actionable therapeutic vulnerability. To cope with stalled replication forks, cells employ a signaling cascade known as the S phase checkpoint to ensure that cells do not progress to mitosis when the DNA is incompletely replicated. CHK1, which is phosphorylated when the checkpoint is activated, is a central node for this checkpoint pathway. We detected more pCHK1-S345 by IF in ecDNA containing tumour cells compared with the corresponding isogenic HSR cells (**Fig. 3a, Extended Fig. 6a**), indicating that transcription replication conflict on ecDNA leads to S phase checkpoint activation in ecDNA-containing tumour cells. In the absence of a functioning checkpoint, cells with highly damaged DNA proceed through the cell cycle, leading to cell death^24^. We therefore hypothesized that ecDNA oncogene amplified tumour cells, due to their intrinsic heightened RS, would be hyper-reliant on CHK1 to manage DNA damage and that disruption of CHK1 could trigger preferential cell death in ecDNA-containing tumour cells.

To test this hypothesis, we used CRISPR to knock out the gene encoding CHK1 in a pair of Hela cell lines with or without *DHFR* amplification on ecDNA. Two different sgRNAs targeting CHK1 induced two- to three-fold higher growth inhibition in ecDNA-positive compared with ecDNA-negative HeLa cells across different time points (**Fig. 3c**). We next inhibited CHK1 pharmacologically using CHIR-124^25^ and found that ecDNA-containing tumour cells were more sensitive to CHK1i than their corresponding isogenic HSR cells, with an IC50 approximately four-fold higher in COLO320HSR compared to COLO320DM cells (**Fig. 3d**). The susceptibility of ecDNA-containing tumour cells to CHK1 inhibition was confirmed with three structurally different CHK1 inhibitors, whereas the CHK2 inhibitor showed no differential inhibitory effect between ecDNA+ and ecDNA- isogenic cell lines (**Extended Fig. 7a-e**). More importantly, inhibition of cell growth by CHK1i was mediated through induction of cell death, as a more rapid and higher degree of cell apoptosis was observed in ecDNA-containing tumour cells treated with CHK1i, as detected by TUNEL (**Fig. 3e**) and PI-Annexin V staining (**Extended Fig. 7f**).

As a master effector of S phase checkpoint, CHK1 activation maintains cell viability by restricting cell cycle progression^24,26^, limiting late replication origin firing to prevent excessive DNA damage accumulation, and protecting stalled replication forks^27,28^. γH2AX IF combined with EdU labeling in COLO320DM and COLO320HSR cells treated with CHK1i showed that CHK1i induced significantly higher DNA damage in COLO320DM compared with COLO320HSR cells, especially in S phase cells as indicated by EdU+ staining, consistent of the function of CHK1 in replication (**Fig. 3f**). Furthermore, inhibition of replication origin firing by CDC7i (XL413), indicated by the decreased EdU-staining intensity (**Extended Fig. 7g**), partially blocked DNA damage induced by CHK1i (**Fig. 3f**), suggesting that CHK1 inhibition leads to extensive replication stress and DNA damage partially through unscheduled replication origin firing.

Taken together, our findings demonstrate that transcription replication conflict, replication stress, and increased baseline DNA damage are common features of ecDNA amplicons and drive activation of the S phase checkpoint. Targeted CHK1 inhibition in ecDNA- positive cells leads to unscheduled replication origin firing and accumulation of DNA damage. Furthermore, the high levels of transcription replication conflict and replication stress drive a selective vulnerability to CHK1 inhibition in ecDNA-positive cells compared to ecDNA- negative cells, raising the possibility for an effective ecDNA-targeted therapy (**Fig. 3g**).

## Oral CHK1i in combination with a pan-FGFRi demonstrates synergistic anti-tumour activity, and inhibits acquired resistance to targeted therapy manifested by ecDNA

Despite convincing preclinical data and preliminary evidence of single agent clinical activity for CHK1 inhibition, there are currently no approved CHK1 inhibitors for any cancer indication. Several limitations of prior CHK1 inhibitors include insufficient potency, potential off-target liabilities (e.g., checkpoint kinase 2 CHK2), and overlapping toxicity in combination with DNA-damaging chemotherapy^29^. To further interrogate the potential of CHK1 inhibition as a treatment for ecDNA-positive cancers, we developed BBI-2779, an orally bioavailable, potent, and selective small molecule inhibitor of CHK1 (**Fig. 4a)**. The potency of BBI-2779 against CHK1 was confirmed *in vitro* using biochemical enzyme inhibition and cellular biomarker assays. The biochemical inhibition IC50 of BBI-2779 against CHK1 was found to be 0.3 nM, while cellular induction of replication stress (as judged by pCHK1-S345, due to CHK1 phosphorylation by upstream kinases) in tumour cells was confirmed to be 3 nM. BBI-2779 has superior biochemical and selective cell growth inhibition compared to other orally bioavailable CHK1 inhibitors tested (IC50 of ecDNA+ CTG proliferation is ∼18-168-fold more potent, **Fig. 4b**). The inhibitor was confirmed to be >160-fold selective for CHK1 over CHK2, suggestive of high pharmacological specificity (**Fig. 4b)**. BBI-2779 also displays excellent bioavailability (%F=71) and good exposure in rodent, allowing for robust CHK1 target coverage after oral administration (**Extended Figs 8a-b**).

Since ecDNA+ oncogene amplified tumour cells harbor elevated intrinsic replication stress and are sensitive to other CHK1 inhibitors (**Fig. 3d**), we hypothesized that they would also be hypersensitive to BBI-2279. Consistent with this notion, BBI-2779 treatment of COLO320DM cells resulted in a significantly greater dose-dependent increase in the expression of the replication stress biomarker pRPA2-S8 compared to COLO320HSR (**Fig. 4c).** COLO320DM cells also showed a greater dose-dependent increase of pCHK1-S345 and γH2AX, as determined by Western blotting (**Fig. 4d)**. In further support of high selectivity, the concentration-dependent induction of replication stress induced by BBI-2779 directly correlated with enhanced cytotoxicity in the COLO320DM cells as compared to COLO320HSR cells, with an approximately 10-fold difference in IC50 between COLO320DM and COLO320HSR cells, demonstrating synthetic lethality in the ecDNA+ context (**Fig. 4e).**

Applying targeted therapy pressure to the protein products of oncogenes amplified on ecDNA induces cancer cells to evade such pressures, either by increasing ecDNA amplification of the dominant oncodriver (**Extended Figs. 9a-d)**, or by ecDNA amplification of a new bypass oncogene^30^. We therefore investigated whether combining targeted therapy with CHK1 inhibition in ecDNA amplified tumour cells provides a synergistic therapeutic effect resulting in cancer cell death and tumour regression. The synergistic antitumour activity and pharmacodynamics of BBI-2779 was evaluated in combination with the pan-FGFR tyrosine kinase inhibitor, infigratinib, in the *FGFR2* amplified ecDNA+ gastric cancer SNU16 xenograft tumour model.

Single agent BBI-2779 or infigratinib resulted in significant tumour growth delay with mean % Tumour Growth Inhibition (ΔTGI) of 64% and 97% compared to the vehicle arm on Day 25 (P < 0.05 and P < 0.0005, respectively) (**Fig. 4f)**. Prolonged treatment of SNU-16 tumour cells *in vitro* and SNU-16 xenograft tumours *in vivo* with infigratinib, resulted in tumour cell stasis for a period of 1-2 weeks, followed by acquired resistance to infigratinib and re-initiation of tumour growth concomitant with increased *FGFR2* amplifications on ecDNA (**Fig. 4g, Extended Figs. 9a-c)**. The lack of robust or sustained anti-tumour activity observed with infigratinib alone is consistent with the absence of compelling clinical efficacy reported for pan-FGFR inhibitors in FGFR1/2/3 amplified settings^31^. Increased *FGFR2* gene amplification correlated with FGFR2 protein levels that likely out-titrate the exposure of infigratinib at its maximally tolerated dose in mice (**Fig. 4h, Extended Fig. 9d)**. The combination of BBI-2779 plus infigratinib resulted in significant TGI from vehicle treated animals (P < 0.0001) with tumour regressions observed over the duration of the study, which was directly correlated with the suppression of further (adaptive) *FGFR2* oncogene copy number amplification on ecDNA, otherwise induced by single agent infigratinib (**Figs. 4f-g**). As expected, both single agent BBI- 2779 and combination of BBI-2779 plus infigratinib treatment resulted in a heightened tumour expression of RS biomarkers pCHK1-S345 and pRPA2-S8 compared to vehicle treated tumours (**Fig. 4h)**. Taken together, these findings demonstrate synergistic anti-tumour activity by combining a selective CHK1 inhibitor with a targeted therapy against the amplified driver oncogene to attenuate ecDNA-mediated resistance. Uncontrolled origin firing caused by selective CHK1 inhibition severely disrupts oncogene expression on hyper-transcribed ecDNA templates, thereby rendering the oncogene addicted tumour cells highly vulnerable to FGFR inhibition.

## Discussion

EcDNA is a pernicious driver of tumour evolution because it is a platform for massive oncogene expression and rapid genome adaptation. Here we show that the transcriptional advantage of ecDNA can be turned on its head to selectively target ecDNA containing tumours. The increased transcription of ecDNA is not limited to the protein-coding oncogene loci, but also extends to multiple noncoding intergenic and antisense regions throughout ecDNAs. The pervasive transcription initiation is consistent with increased chromatin accessibility and promiscuous enhancer-promoter contacts on ecDNA^9,10^. Thus, rampant ecDNA transcription comes at the cost of increased transcription replication conflict that cancer cells must manage. DNA damage has been previously associated with ecDNA containing cancers principally as a source of ecDNA generation^8^. Our results show that, once formed, ecDNAs become themselves a major driver of DNA damage. The RNA transcription and DNA replication machineries are two processive holoenzymes that both run along DNA; they must take turns or risk collision. EcDNA containing cancer cells are balanced on the edge of DNA damage catastrophe. We find that ecDNAs are constantly breaking due to transcription replication conflict, and cancer cells become heavily reliant on the S-phase checkpoint kinase CHK1 to limit origin firing. The alternative for the cancer cell is to limit ecDNA transcription and lose oncogene overexpression, undermining their unique oncogenic and adaptive growth advantage.

We tested the concept that enhancing transcription replication conflict will cause ecDNA containing tumour cells to self-destruct. Inhibition of CHK1 substantially increases ecDNA damage during DNA replication and leads to preferential killing of ecDNA containing cancer cells. There are currently no approved CHK1 inhibitors for use in cancer patients. Despite convincing preclinical data and preliminary evidence of single agent clinical activity for CHK1 inhibition, a predictive biomarker(s) and an optimal clinical development strategy have been lacking. Furthermore, a major challenge to the successful clinical development of CHK1 inhibitors has been the lack of reliable methods to identify high-RS tumours that are predicted to be hypersensitive to CHK1 inhibition^32–39^. Long durability of CHK1 inhibition in vivo is likely required to exploit unscheduled DNA replication to ensure cancer cell death. The results presented here suggest a promising strategy for a next generation CHK1 inhibitor to target ecDNA containing cancers. Notably, CHK1 inhibition showed synergy with a targeted therapy blocking the ecDNA oncogene-encoded product and prevented the adaptive elevation of ecDNA copy number that previously foiled monotherapy targeting ecDNA protein products. Previous successes in cancer therapy have exploited the synthetic lethality of deficiencies, for example PARP inhibition in BRCA2 deficient cancer cells^40^. Our work demonstrates the feasibility of a synthetic lethality of excess to turn the molecular advantages of ecDNA in cancer against itself.

## Author Contributions

H.Y.C. and P.S.M. conceived the study. J.T., E.J.C., I.T-L.W. conducted studies of ecDNA replication stress, CHK1 activation, and synthetic lethality. N.E.W, G.W., G.M., R.L. conducted genomic studies of ecDNA transcription, ssDNA accumulation, and response to triptolide.

W.J.G. and V.B. supported analyses. S.C. mapped ecDNA replication speed. P.H., S.J.B. conducted comet assays of ecDNA breakage. E.T., R.H., J.P., A.S., S.T.M., T.P., S.K., C.A.H. conducted CHK1 knockout experiments and designed, synthesized, and evaluated BBI-2779. J.T., N.E.W., G.W., S.C., H.Y.C. wrote the manuscript with input from all coauthors. C.A.H., P.S.M., H.Y.C. supervised their respective teams for the study.

## Acknowledgements

We thank members of the Chang and Mischel labs for discussion. This work was delivered as part of the eDyNAmiC team supported by the Cancer Grand Challenges partnership funded by Cancer Research UK (CGCATF-2021/100012 and CGCATF-2021/100025) and the National Cancer Institute (OT2CA278688 and OT2CA278635) to H.Y.C., P.S.M., and V.B. This project was supported by NIH RM1-HG007735 (H.Y.C., W.J.G., C.C.). Confocal microscopy was performed on instruments in the Stanford CSIF core facility (supported by RRID:SCR_017787). FACS analysis was performed on instruments in Stanford Shared FACS Facility. H.Y.C. is an Investigator of the Howard Hughes Medical Institute.

## Data and Material Availability

All genomic data will be publicly available in Gene Expression Omnibus (https://www.ncbi.nlm.nih.gov/geo/) upon publication. BBI-2779 is available upon request to C.A.H. at Boundless Bio.

## Extended Data Figures

**Extended Data Figure 1.**
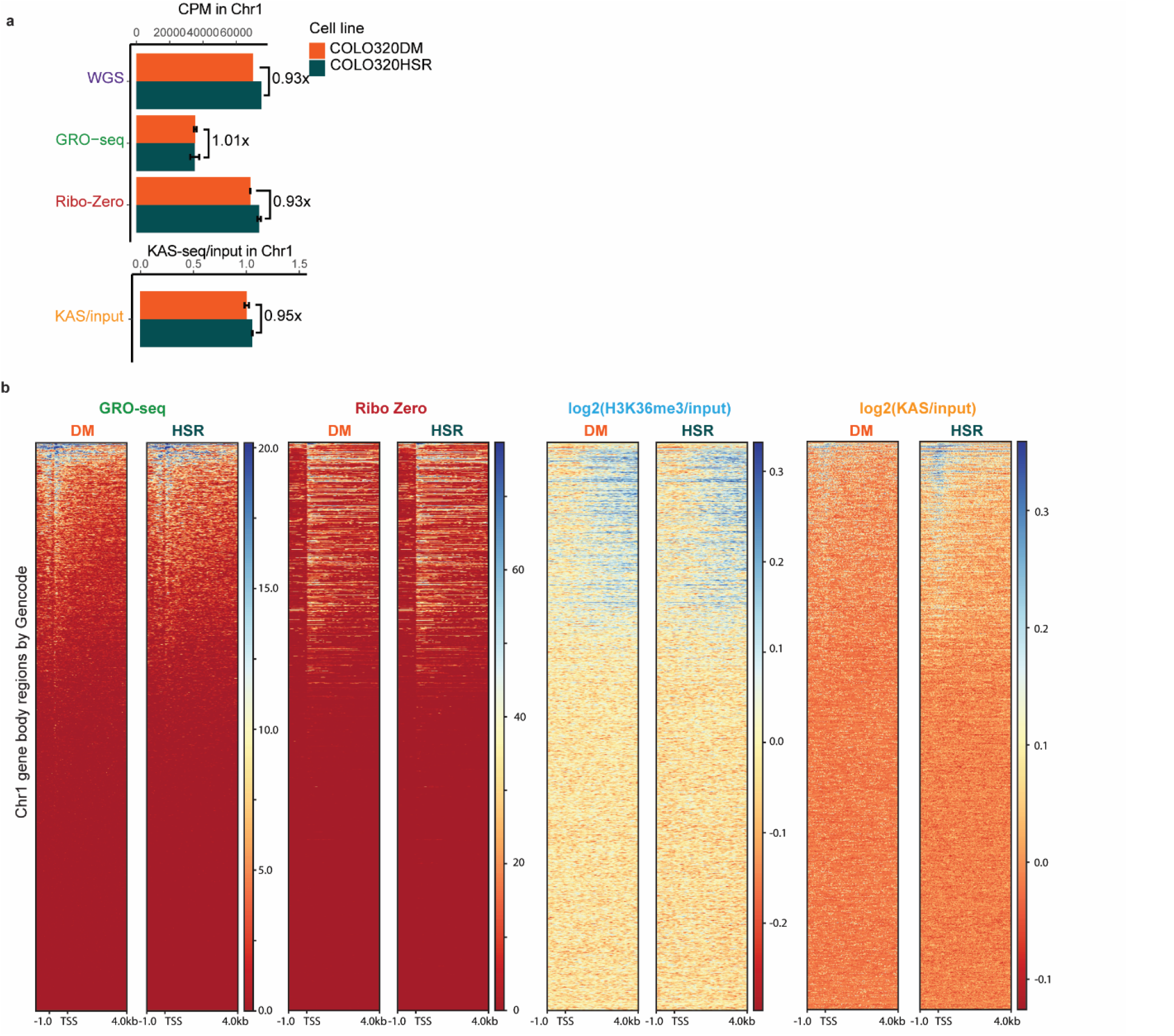
Similar levels of transcription and ssDNA accumulation on normal chromosome 1 of COLO320 cell lines. **(a)** Read density of genomic assays in COLO320DM and COLO320HSR in total counts per million (CPM) on chromosome 1, which is outside of ecDNA intervals. KAS-seq read density is shown as (CPM) of the KAS-seq relative to CPM of the input. For GRO-seq, Ribo-Zero, and KAS-seq the mean of two biological replicates is shown, error bars indicate standard deviation. For WGS, a single representative replicate is used^11^. **(b)** Metagene heatmap plot visualization of GRO-seq, Ribo Zero RNA-seq, log2 of input-normalized H3K36me3 ChIP-seq, and log2 of input-normalized coverage of KAS-seq within chromosome 1. All plots are anchored at the GRO-seq TSS as identified by HOMER. One representative biological replicate for each condition is visualized.

**Extended Data Figure 2.**
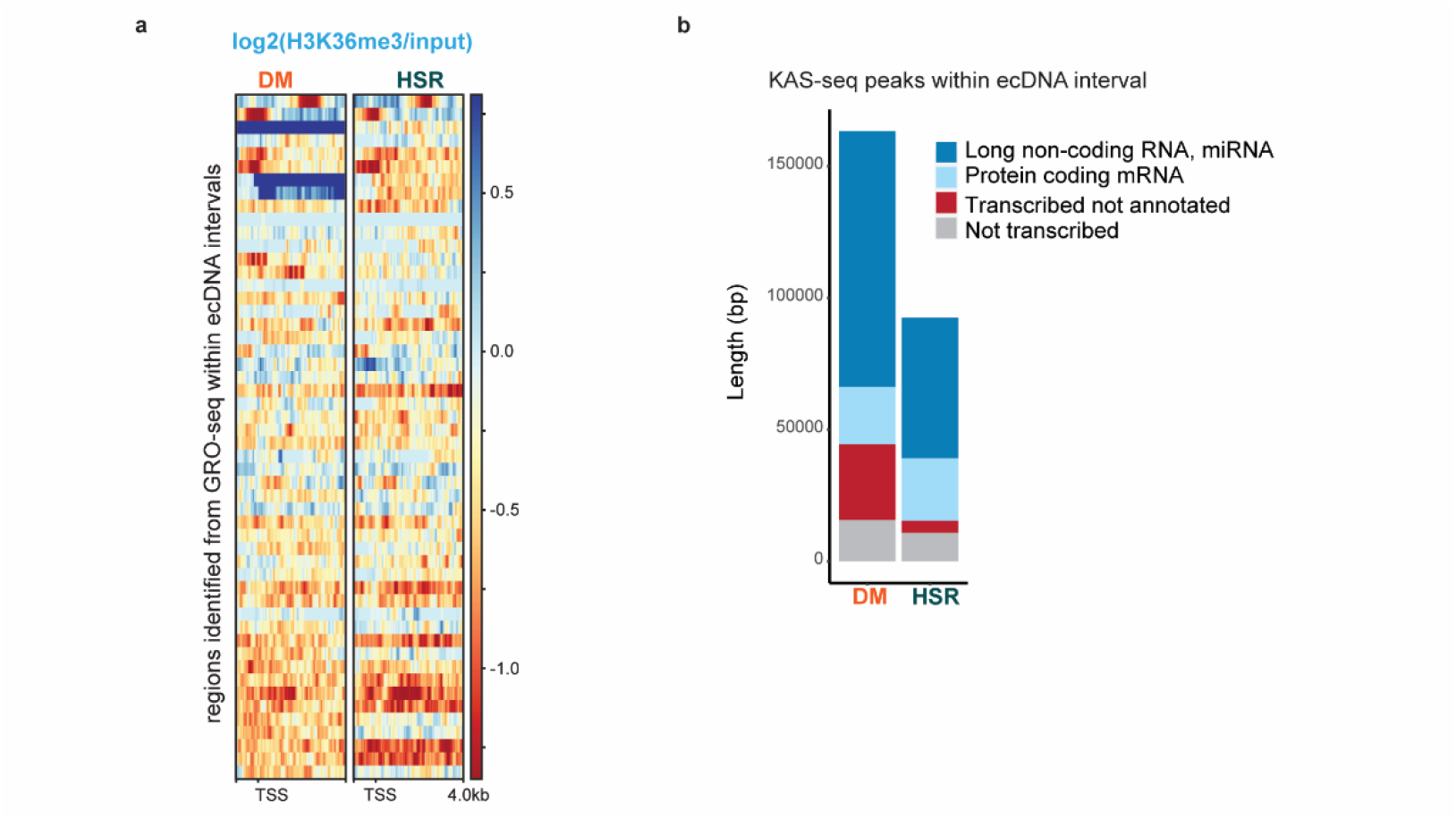
H3K36me3 and KAS-seq signals within the ecDNA interval of COLO320 cell lines. **(a)** Metagene heatmap plot visualization of log2 of input-normalized H3K36me3 ChIP-seq within the ecDNA interval. Plots are anchored at the GRO-seq TSS as identified by HOMER. One representative biological replicate for each cell line is visualized. **(b)** Accumulative bar plots of length distributions of all KAS-seq peaks identified within the ecDNA interval classified by GRO-seq transcription status and Gencode v43 annotation. Two biological replicates were used per cell line.

**Extended Data Figure 3.**
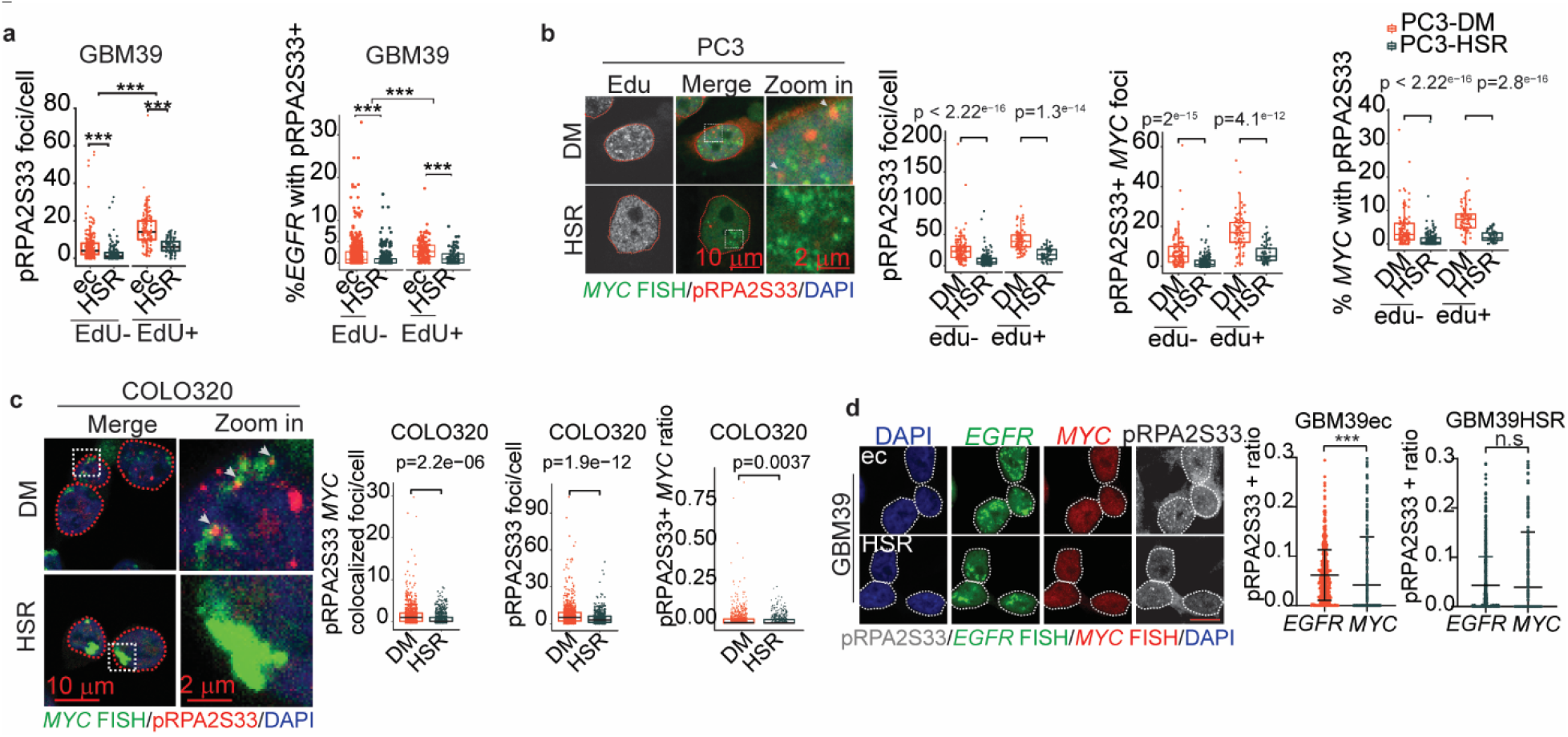
Replication stress on ecDNA with different amplification sequence in different tumour cells. (**a**) Quantification of images in Fig 2e, left: total pRPA2-S33 foci/cell; right: % of *EGFR* co- localized with pRPA2-S33. (Box plot parameters same as in Fig 1b, Two-tailed Mann-Whitney test, EdU- group: GBM39ec, n=244, GBM39HSR, n=143; EdU+ group: GBM39ec, n=104, GBM39HSR, n=72). (**b**) pRPA2-S33 immunofluorescence combined with *MYC* FISH staining in PC3-DM and PC3-HSR cells, with EdU added 30 min before fixation. Left: representative images. 2nd left: pRPA2-S33 foci number. 2nd right: Quantification of *MYC* foci number colocalized with pRPA2-S33. Right: Quantification of percentage of *MYC* co-localized with pRPA2-S33. (Box plot parameters were the same as in Fig 1b, two-tailed Mann-Whitney test, EdU+ group: PC3-DM, n=81, PC3-HSR, n=58; EdU- group: PC3-DM, n=128, PC3-HSR, n=184). (**c**) pRPA2-S33 immunofluorescence combined with *MYC* FISH staining in COLO320DM and COLO320HSR cells. Left: representative images. 2nd left: pRPA2-S33 colocalized with *MYC* foci number. 2nd right: pRPA2-S33 foci number. Right: ratio of *MYC* that colocalized with pRPA2-S33. (Box plot parameters were the same as in Fig 1b, two-tailed students’ t test, COLO320DM, n=974; COLO320HSR, n=495). (**d**) Left, representative images of pRPA2-S33 immunofluorescence staining combined with *EGFR* and *MYC* FISH staining. Middle and right, proportion of *EGFR* and *MYC* locus co-localized with pRPA2-S33 in GBM39ec cells (middle panel) and GBM39HSR cells (right panel). (mean± SD, two-tailed students’ t test, GBM39ec, n=493; GBM39HSR, n=614) (* represents p<0.05, ** represents p<0.01, *** represents p<0.001, scale bar represents 10 µm or as otherwise specified in the image)

**Extended Data Figure 4.**
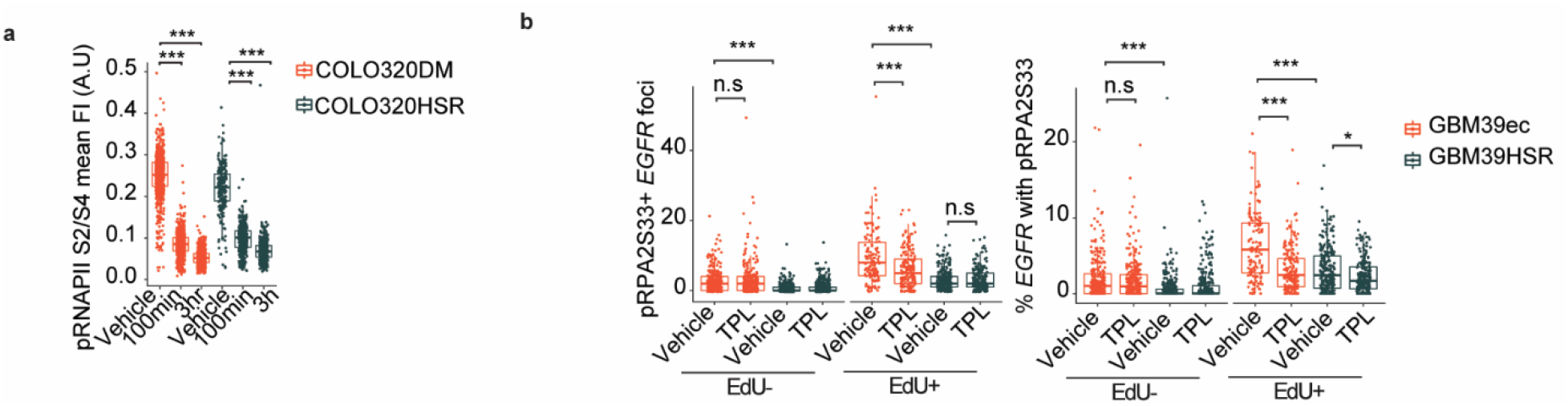
Transcription replication conflict drives replication stress in ecDNA containing tumour cells. (**a**) Mean pRNAPolII S2/S4 fluorescence intensity (arbitrary units) was measured in datasets shown in Fig 2h. (Box plot parameters were the same as in Fig 1b, two-tailed Wilcoxon test, COLO320DM group: vehicle, n=354, 100 min, n=350, 3 h, n= 269; COLO320HSR group: vehicle, n=130, 100 min, n= 185, 3 h, n= 161). (**b**) Quantification of dataset shown in Fig 2i. left, pRPA2-S33 and *EGFR* colocalized foci number; right, percentage of *EGFR* colocalized with pRPA2-S33. (Box plot parameters were the same as in Fig 1b, two-tailed Wilcoxon test, sample size from left to right: n= 280, 256, 332, 348, 138, 189, 253, 208) (* represents p<0.05, ** represents p<0.01, *** represents p<0.001)

**Extended Data Figure 5.**
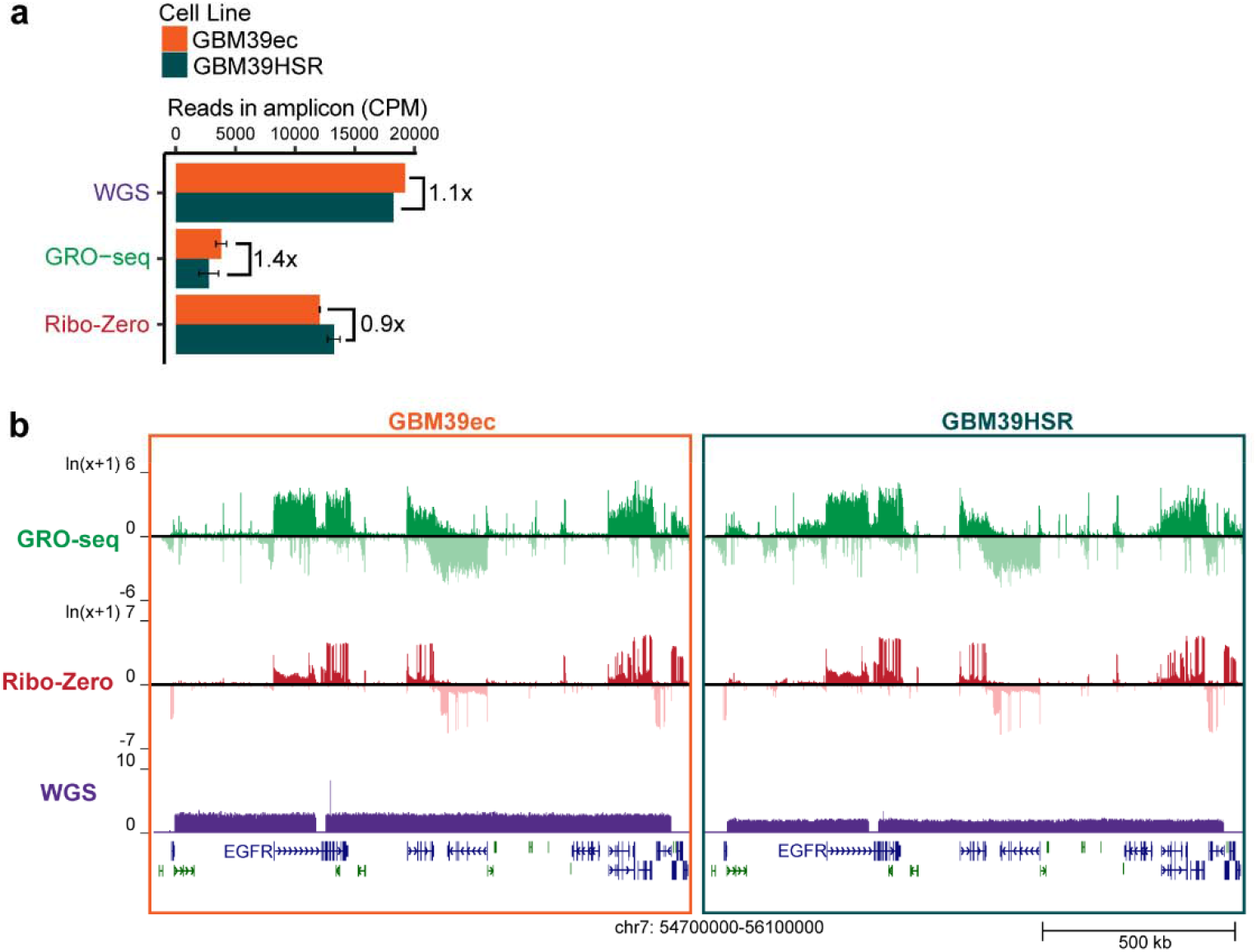
Transcription within the ecDNA interval of GBM39 cell lines. **(a)** Read density of genomic assays in GBM39ec and GBM39HSR within the ecDNA interval in total counts per million (CPM). For GRO-seq and Ribo-Zero the means of two biological replicates are shown, error bars indicate standard deviation. For WGS, a single representative replicate is used^11^. **(b)** Genome tracks highlighting the GBM39 ecDNA interval. One representative biological replication for each condition is visualized.

**Extended Data Figure 6.**
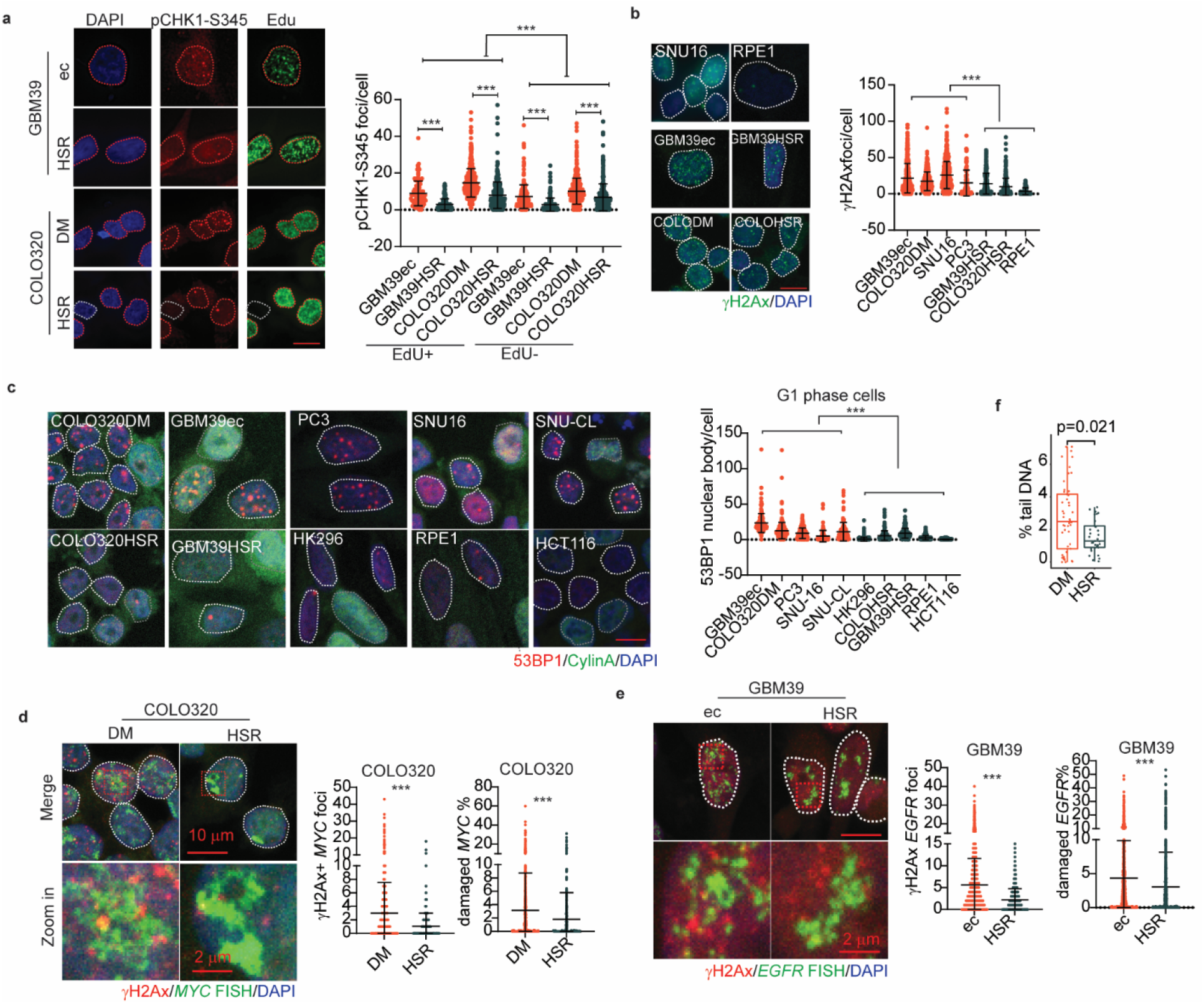
Replication stress induces DNA lesions and activates S phase check point in ecDNA containing tumour cells. (**a**) pCHK1-S345 staining in 2 isogenic cell line pairs, GBM39ec, GBM39HSR and COLO320DM, COLO320HSR, with EdU added 30min before fixing samples. Left, representative images, red dotted lines mark EdU+ nuclei and white dotted lines mark EdU- nuclei; right, quantification of pCHK1 foci number. (mean± SD, two tailed Mann-Whitney test, sample size from left to right, n= 97, 192, 566, 339, 267, 252, 610). (**b**) γH2AX staining in cell line panels with or without ecDNA. Left, representative image; right, quantification of γH2AX foci number per cell. (mean± SD, two tailed Mann-Whitney test, sample size from left to right, n= 402, 362, 499, 101, 388, 418, 80). For COLO320DM and COLO320HSR the same images are shown in Figure 3a; they are used here to compare against multiple cell lines. (**c**) 53BP1 combined with CyclinA staining in cell line panels with or without ecDNA. Left, representative image. White dotted lines mark G1 phase cells which are Cyclin A negative nuclei and grey dotted lines mark Cyclin A positive nuclei. Right, quantification of 53BP1 in G1 phase cells which were CyclinA negative cells. (mean± SD, two tailed Mann-Whitney test, sample size from left to right, n= 180, 277, 240, 146, 126, 211, 188, 163, 221, 494). For COLO320DM and COLO320HSR the same images are shown in Figure 3a; they are used here to compare against multiple cell lines. (**d**) γH2AX IF combined with *MYC* FISH staining in COLO320DM and COLO320HSR cells. Left: representative images. Middle, quantification of γH2AX colocalized with *MYC* foci number, right, quantification of percentage of *MYC* colocalized with γH2AX. (mean± SD, two tailed Mann-Whitney test, sample size, COLO320DM, n=804; COLO320HSR, n=411). (e) γH2AX IF combined with *EGFR* FISH staining in GBM39ec and GBM39HSR cells. Left, representative image. Middle, quantification of γH2AX colocalized with *EGFR* foci number, right, quantification of percentage of *EGFR* colocalized with γH2AX. (mean± SD, two tailed Mann-Whitney test, sample size, GBM39ec, n=1638; GBM39HSR, n=1863). (**f**) Quantification of percentage of tail DNA content with the dataset shown in Fig 3b. (Box plot parameters were same as in Fig 1b, two-tailed Wilcoxon test, COLO320DM, n= 49, COLO320HSR, n= 33). (* represents p<0.05, ** represents p<0.01, *** represents p<0.001, scale bar represents 10 µm or indicated in the images)

**Extended Data Figure 7.**
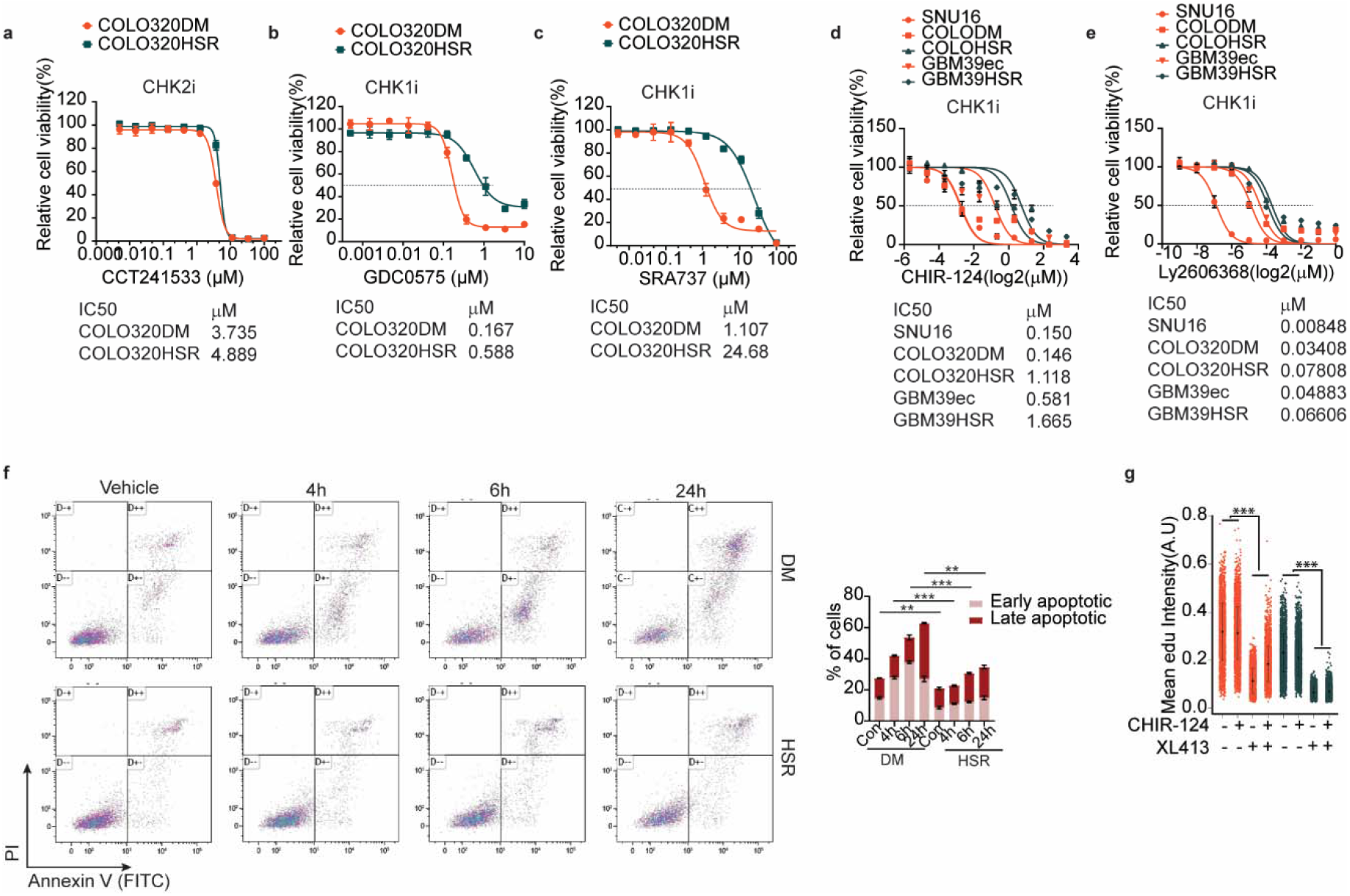
ecDNA containing tumour cells are sensitive to targeted CHK1 inhibition. (**a-e**) Cell viability curve of COLO320DM, COLO320HSR, GBM39ec, GBM39HSR and SNU16 in response to different chemicals targeting CHK1 or CHK2. a. CHK2i, CCT241533; b. CHK1i, GDC0575; c. CHK1i, SRA737; d, CHK1i, CHIR-124; e, CHK1i, Ly2606368. Half maximal inhibitory concentration (IC50) for each inhibitor in individual cell lines were listed on the bottom. (sample size in a-c, n=2; d-e, n=4) (**f**) FACS analysis of Annexin V staining in COLO320DM and COLO320HSR cells subjected to 1 µM CHIR-124 for indicated time. Left, gating setting and representative plots, right, % of early apoptotic cells: Annexin V + and PI-; late apoptotic cells: Annexin V + and PI +. (mean± SD, P values quantified by two-tailed students’ t test, n=2.) (**g**) Quantification of mean EdU intensity (arbitrary units) in dataset shown in Fig 3f. (mean± SD, P values quantified by two-tailed students’ t test. Sample size from left to right: n= 419, 284, 1085, 596, 209, 242) (* represents p<0.05, ** represents p<0.01, *** represents p<0.001)

**Extended Data Figure 8.**
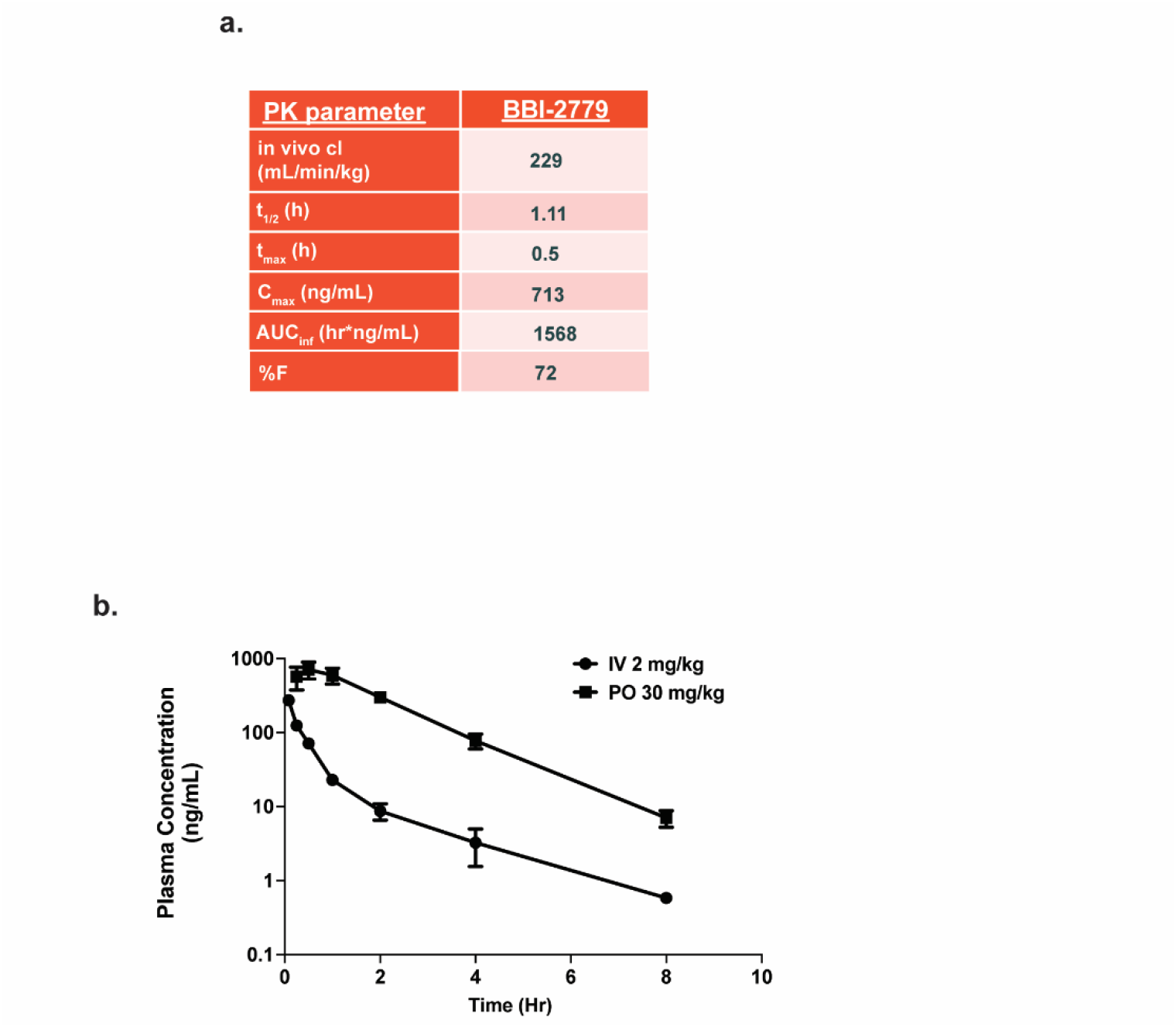
BBI-2779 is well tolerated and illustrate optimal PK exposure in mice. (**a**) Pharmacokinetic parameters of BBI-2779. Oral bioavailability was determined in fasted male CD-1 mice dosed at 30 mg kg-1 (n = 3). (**b**) Plasma concentration time curve of BBI-2779 in mouse either administered intravenously (IV) at 2mg kg-1 or orally (PO) at 30 mg/kg-1. Data are mean ± s.e.m., n = 3 per group.

**Extended Data Figure 9.**
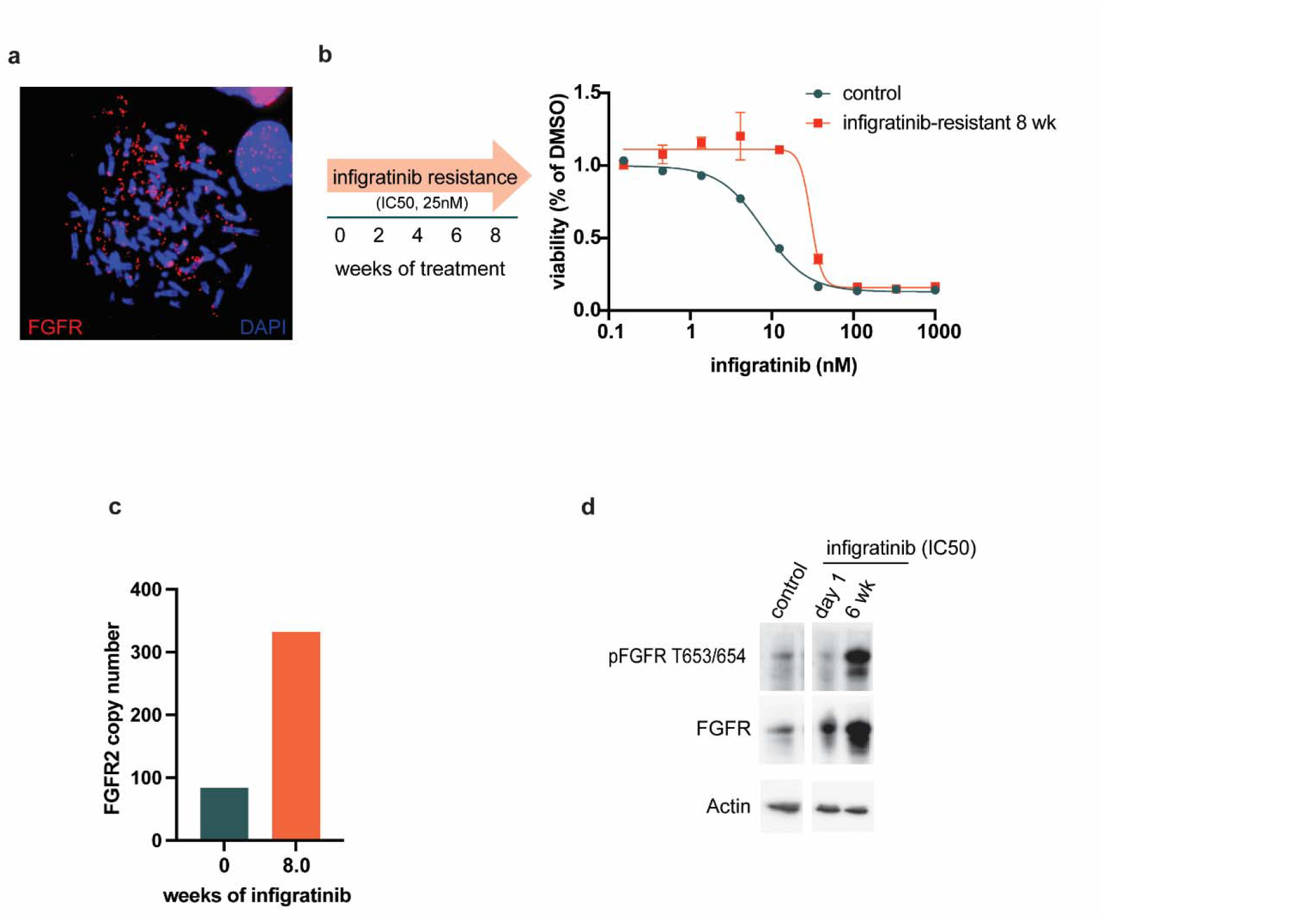
Targeted therapeutic resistance shaped by intracellular ecDNA - driven oncogene amplification. (**a**) *FGFR2* (red) FISH imaging of cells in metaphase demonstrate amplification of *FGFR2* oncogene on ecDNA in SNU16 cells. Nuclear staining is illustrated using DAPI (blue). (**b**) Timeline of experimental overview. After 8 weeks of infigratinib treatment (EC50 dose of 25 nM), cells were assessed for infigratinib resistance. A 3-day Cell Titer Glo reveals resistance in SNU16 cells treated with infigratinib. (**c**) qPCR based quantification of FGFR2 oncogene numbers after 8 weeks of infigratinib treatment showing SNU16 cells resistant to infigratinib with significant increase in FGFR2 target selection/amplification. (**d**) Western blotting illustrating enhanced expression of FGFR signaling pathways involved in therapeutic resistance.

## Methods

### Antibodies and reagents

Antibody: H3K36me3 (abcam ab9050), γH2AX (Millipore, 05-636 for immunofluorescence), γH2AX (Cell Signaling Technology,, CST9718 for western blot), pRPA2S33 (Novus Biological, NB100-544), pCHK1S345 (Invitrogen, PA5-34625), 53BP1 (Novus Biological, NB100-304), cyclin A (BD Bioscience, #611268), pRNAPII S2/S4 (Abcam, ab252855), pCHK1-S345 (Cell Signaling Technology, CST2348), CHK1 (Abcam, ab32531), pRPA32/RPA2-Ser8 (Cell Signaling Technology, 54762S), Vinculin (Cell Signaling Technology, CST13901), pFGFR2- Tyr653/654 (Cell Signaling Technology, CST3476S), FGFR2 (Cell Signaling Technology, CST11835S).

Chemicals: CHIR-124 (Selleck chem, Catalog No. S2683); XL413 (Selleck chem, Catalog No. S7547); Triptolide (Millipore, 645900-5MG).

### Cell culture

GBM39ec, GBM39HSR and HK296 were patient derived neurosphere cell lines and were established as previously described^2,7^. All the other cell lines were purchased from ATCC. Human prostate cancer cell line PC3 DM, PC3 HSR; colorectal cancer cell line COLO320DM, COLO320HSR; gastric cancer cell line SNU16; lung cancer cell line PC9 and hTERT-immortalized retinal pigment epithelial cell line RPE1 were cultured in 4.5 g/L glucose formulated Dulbecco’s Modified Eagle’s Medium (DMEM, Corning) supplemented with 10% FBS (Gibco). For GRO-seq and ChIP-seq, COLO320DM and COLO320HSR were grown in Roswell Park Memorial Institute (RPMI) 1640 with Glutamax (Gibco) with 10% FBS. GBM39ec, GBM39HSR and HK296 cell lines were cultured in DMEM/F12 (Gibco, 11320-033) supplemented with 1 × B27 (Gibco, 17504-01), 20 ng/ml EGF (Sigma, E9644), 20 ng/ml FGF (Peprotech, AF-100-18B) and 1-5 µg/ml Heparin (Sigma, H3149) and 1 × Glutamax (Gibco, 35050-061). GBM39 cells used in sequencing assays were cultured without additional Glutamax. All the cells were maintained at 37 ° C in a humidified incubator with 5% CO_2_. Cell lines routinely tested negative for mycoplasma contamination.

### GRO-seq

COLO320DM and COLO320HSR RNA was prepared by washing cells with ice cold PBS, then adding ice-cold LB (10mM Tris-HCl pH 7.4, 2mM MgCl2, 3mM CaCl2, 0.5% IGEPAL-CA630, 10% glycerol, 1mM DTT, protease inhibitors (Roche 11836170001), RNAse inhibitor (Ambion AM2696)) and scraping cells into a 15ml conical tube. Cells were spun at 1000g for 10min at 4 °C. Supernatant was removed and pellet was thoroughly resuspended in 1ml LB using a wide bore tip. An additional 9ml LB was added and then cells were spun at 1000g for 10min at 4 °C. Cells were resuspended in LB again and spun down. Pellets were resuspended in ice-cold freezing buffer (50mM Tris-HCl pH 8.3, 5mM MgCl2, 40% glycerol, 0.1mM EDTA, 0.2ul RNAse inhibitor per ml of freezing buffer and spun at 2000g for 2min at 4 °C. Nuclei were resuspended in 100μl freezing buffer/5 million cells. A nuclear run-on master mixed was prepared (10mM Tris-HCl pH 8.0, 5mM MgCl2, 1mM DTT, 300mM KCl, 0.5mM ATP, 0.5mM GTP, 0.003mM CTP (unlabeled rNTPs from Roche 11277057001), 0.5mM Bromo-UTP (Sigma B7166), 1% Na-laurylsarcosine, 1μl RNase inhibitor/100μl) and preheated to 30°C. An equal volume of master mix was added to aliquoted nuclei (5 million nuclei per replicate) and incubated at 30°C for 5 minutes with gentle shaking. DNAse digestion was performed using RQ1 DNase I and RQ1 buffer (Promega M610A) for 30min at 37°C, the reaction was stopped with the addition of stop buffer to a final concentration of 10mM Tris-HCl pH 7.4, 1% SDS, 5mM EDTA, 1mg/ml Proteinase K. Samples were incubated for 1hr at 55°C. NaCl was added to final concentration of 225mM. Two phenolchloroform extractions were done followed by one extraction with chloroform. RNA was precipitated in 75% EtOH with 1μl glycoblue (Ambion 9516) overnight at -20 °C.

For GBM39ec and GBMHSR, cells were washed with ice-cold PBS and then spun for 5 min at 500g, 4C. cells were then resuspended in ice-cold 10ml swelling buffer (10mM Tris-HCl pH 7.5, 2mM MgCl2, 3mM CaCl2, protease inhibitor, RNAse inhibitor) and inclubated on ice for 5 minutes. Cell were spun at 400g for 10min at 4°C and resuspended in 10ml ice-cold glycerol swelling buffer (0.9x swelling buffer, 10% glycerol). 10ml ice-cold lysis buffer (glycerol swelling buffer,1% IGEPAL-CA630) was slowly added while agitating the tube. Samples were incubated on ice for 5 minutes, then another 25ml lysis buffer was added and samples were spun for 5min at 600g, 4°C. Samples were resuspended in ice-cold freezing buffer (50mM Tris-HCl pH 8.0, 5mM MgCl2, 40% glycerol, 0.1mM EDTA, RNAse inhibitor) and spun at 900g for 6min 4°C. An equal volume of pre-warmed nuclear run-on master mix was added to aliquoted nuclei (10 million nuclei per replicate) and incubated at 30°C for 7 minutes with gentle shaking. Samples were then mixed thoroughly with 600μl Trizol LS and incubated at room temperature for 5 min. 160μl chloroform was added to each sample, shaken vigorously, then incubated and room temperature for 3 min, centrifuged at 12,000g at 4 °C for 30min. NaCl was added to the aqueous phase to a final concentration of 300mM and RNA was precipitated in 75% EtOH with 1ul glycoblue overnight at -20 °C.

For all cell types, after overnight RNA precipitation, RNA was spun for 20 min at 21130g, 4°C. RNA pellets were washed in fresh 75% EtOH, briefly air-dried, and then resuspended in 20ul water. Base hydrolysis was performed using 5ul 1N NaOH for 10 minutes and then neutralized with 25ul 1M Tris-HCl pH 6.8. Buffer exchange was performed using P30 Micro columns (Biorad 732-6250), then treated with RQ1 DNase I and RQ1 buffer and incubated at 37°C (10min for COLO320 and 30min for GBM39). Buffer exchange was performed again. Samples were treated with 3ul T4 PNK (New England Biolabs M0201), 1x PNK buffer, 2ul 10mM ATP, 2ul RNase inhibitor and incubated for 1hr at 37C. Another 2 ul pNK was added per sample and incubation was continued for 30-60 minutes. RNA decapping was performed by adding ammonium chloride (final concentration 50mM), poloaxamer 188 (final concentration 0.1%), 2ul mRNA decapping enzyme (New England Biolabs M0608S), and 1ul RNase inhibitor and incubating at 37°C for 30 minutes. EDTA was then added to final concentration of 25mM and samples were incubated at 75C for 5 minutes. Samples were then incubated on ice for at least 2 minutes. Sample volume was then brought to 100ul with binding buffer (0.25X SSPE, 1mM EDTA, 0.05% Tween-20, 37.5mM NaCl, RNAse inhibitor). During T4 PNK treatment, 60ul anti-BrdU agarose beads (Santa Cruz Biotechnology sc-32323ac) per sample were equilibrated in 500ul binding buffer by rotating for 5min at room temperature, spun, and washed again in binding buffer. Beads were then blocked in blocking buffer (1x binding buffer, 0.1% polyvinylpyrrolidone, 1ug/ml ultrapure BSA, RNAse inhibitor) by rotating for 1hr at room temperature. Beads were then washed twice in binding buffer resuspended in 400ul binding buffer. Decapped RNA was then added to the blocked beads and rotated for 1 hour and room temperature. Beads were then washed once in binding buffer, once in low salt buffer (0.2X SSPE, 1mM EDTA, 0.05% Tween-20, RNAse inhibitor), once in high salt buffer (0.2X SSPE, 1mM EDTA, 0.05% Tween-20, 137.5mM NaCl, RNAse inhibitor) with 3 minutes of rotation, and twice in TET buffer (10mM Tris-HCl pH 8.0, 1mM EDTA, 0.05% Tween-20, RNAse inhibitor). All spins with agarose beads were performed for 2 minutes at 1000g at room temperature and all washes were performed in 500ul buffer rotating for 5 minutes at room temperature unless otherwise noted. Samples were then eluted in elution buffer (50mM Tris-HCl pH 7.5, 20mM DTT, 1mM EDTA, 150mM NaCl, 0.1% SDS, RNase inhibitor) pre-warmed to 42°C, four 10 minute elutions were performed at 42C with periodic vortexing. The eluates for each replicate were pooled and RNA was then purified by phenol-chloroform and chloroform with EtOH precipitation (COLO320) or by column purification using New England Biolabs Monarch RNA cleanup kit T2030 (GBM39). Sequencing libraries were prepared using the NEBNext small RNA library prep kit (New England Biolabs E7330) and sequenced by Novaseq PE150. The sequence data were mapped to human reference genome (hg38) using STAR, version 2.7.10b^17^. HOMER (v4.11.1) was used for de novo transcript identification on each strand separately using the default GRO-seq setting. Reads with MAPQ values less than 10 were filtered using SAMtools (v1.8). Duplicate reads were removed using picard-tools. GRO-seq signal was converted to the bigwig format for visualization using deepTools bamCoverage^18^ (v3.3.1) with the following parameters: --binSize 10 --normalizeUsing CPM -- effectiveGenomeSize 3209286105 --exactScaling.

### Total RNA library preparation

Total RNA from each sample was isolated with Quick-RNA Miniprep Kit (Zymo Research, R1054) with input of 1-2 million cells. RNA libraries were constructed using TruSeq Stranded Total RNA Library Prep Kit with Ribo-Zero (#20020596, Illumina). Nextseq 550 sequencing system (Illumina) produced 20-30 million of ×2, 75 base pair, paired-end reads per sample. The sequence data were mapped to human reference genome (hg38) using STAR, version 2.7.10b^17^. Reads with MAPQ values less than 10 and PCR duplicates were filtered using SAMtools (v1.8). Ribo-Zero signal was converted to the bigwig format for visualization using deepTools bamCoverage^18^ (v3.3.1) with the following parameters: --binSize 10 --normalizeUsing CPM -- effectiveGenomeSize 3209286105 --exactScaling.

### KAS-seq library preparation

KAS-seq experiments were carried out as previously described^12^ with modifications^13^. Briefly, cell culture media was supplemented with 5 mM N_3_-kethoxal (final concentration), and cells were incubated for 10 minutes at 37^◦^C in a 6-well plate. Genomic DNA was then extracted using the Monarch gDNA Purification Kit (NEB T3010S) following the standard protocol but with elution using 50 µL 25 mM K_3_BO_3_ at pH 7.0. Click reaction was carried out by mixing 87.5 µL purified DNA, 2.5 µL 20 mM DBCO-PEG4-biotin (DMSO solution, Sigma 760749), and 10 *µ*L 10× PBS in a final volume of 100 µL. The reaction was then incubated at 37^◦^C for 90 minutes.

DNA was purified using AMPure XP beads by adding 50 µL beads 100 µL reaction, washing beads on a magnetic stand twice with 80% EtOH, and eluting in 130 µL 25mM K_3_BO_3_. Purified DNA was then sheared using a Covaris E220 instrument down to ∼200-400 bp size. Pulldown of biotin-labeled DNA was initiated by separating 10 µL of 10 mg/mL Dynabeads MyOne Streptavidin T1 beads (Life Technologies, 65602) on a magnetic stand, then washing with 180 µL of 1× TWB (Tween Washing Buffer; 5 mM Tris-HCl pH 7.5; 0.5 mM EDTA; 1 M NaCl; 0.05% Tween 20). Beads were then resuspended in 300 µL of 2× Binding Buffer (10 mM Tris-HCl pH 7.5, 1 mM EDTA; 2 M NaCl), sonicated DNA was added (diluted to a final volume of 300 µL if necessary), and beads were incubated for ≥15 minutes at room temperature on a rotator. Beads were separated on a magnetic stand and washed with 300 µL of 1× TWB and heated at 55^◦^C in a Thermomixer with shaking at 1,000 rpm for 2 minutes. The supernatant was removed on a magnetic stand, and the TWB wash and 55^◦^C incubation were repeated.

Libraries were prepared on beads using the NEBNext Ultra II DNA Library Prep Kit (NEB, #E7645). First, end repair was carried out by incubating beads for 30 minutes at 20^◦^C in a Thermomixer with shaking at 1,000 rpm in 50 *µ*L 1× EB buffer plus 3 µL NEB Ultra End Repair Enzyme and 7 µL NEB Ultra End Repair Enzyme. This was followed by incubation at 65^◦^C for 30 minutes. Second, adapters were ligated by adding 2.5 µL NEB Adapter, 1 µL Ligation Enhancer, and 30 µL Blunt Ligation mix, incubating at 20^◦^C for 20 minutes, then adding 3 µL USER enzyme, and incubating at 37^◦^C for 15 minutes (in a Thermomixer, with shaking at 1,000 rpm). Beads were separated on a magnetic stand, and washed with 180 µL TWB for 2 minutes at 55^◦^C and 1000 rpm in a Thermomixer. After magnetic separation, beads were washed in 100 µL 0.1 × TE buffer, resuspended in 15 µL 0.1 × TE buffer, and heated at 98^◦^C for 10 minutes. PCR was carried out by adding 5 µL of each of the i5 and i7 NEB Next sequencing adapters together with 25 µL 2× NEB Ultra PCR Mater Mix, with a 98^◦^C incubation for 30 seconds and 15 cycles of 98^◦^C for 10 seconds, 65^◦^C for 30 seconds, and 72^◦^C for 1 minute, followed by incubation at 72^◦^C for 5 minutes. Beads were separated on a magnetic stand, and the supernatant was cleaned up using 1.8× AMPure XP beads.

Libraries were sequenced in a paired-end format on an Illumina NextSeq instrument using NextSeq 550 high-output kits (2×36 cycles). The sequence data were mapped to the hg38 assembly of the human genome using Bowtie^19,20^ with the following settings: -v 2-k 2-m 1--best--strata-X 1000. Duplicate reads were removed using picard-tools (version 1.99). MACS2^21^ (v.2.1.1) was used for peak-calling with the following parameters: --broad -g hs --broad-cutoff 0.01 -q 0.01. Browser tracks are generated after normalizing to input using bamCompare default setting.

### ChIP-seq library preparation

Three million cells per replicate were fixed in 1% formaldehyde for 15 min at room temperature with rotation and then quenched with 0.125 M glycine for 10 min at room temperature with rotation. Fixed cells were pelleted at 1300*g* for 5 min at 4 °C and washed twice with cold PBS before storing at −80 °C. Membrane lysis was performed in 5 ml LB1 (50 mM HEPES pH 7.5, 140 mM NaCl, 1 mM EDTA, 10% glycerol, 0.5% 1% IPEGAL-CA630, 0.25% Triton X-100, and Roche protease inhibitors 11836170001) for 10 min at 4 °C with rotation. Nuclei were pelleted at 1,400*g* for 5 min at 4 °C and lysed in 5 ml LB2 (10 mM Tris-Cl pH 8.0, 200 mM NaCl, 1 mM EDTA, 0.5 mM EGTA, Roche protease inhibitors) for 10 min at room temperature with rotation. Chromatin was pelleted at 1,400*g* for 5 min at 4 °C and resuspended in 1 ml of TE buffer + 0.1% SDS before sonication on a Covaris E220 with the following settings (140 W, 10% duly, 200 cycles/burst, 600 sec/sample). Samples were clarified by spinning at 16,000*g* for 10 min at 4 °C. Supernatant was transferred to a new tube and diluted with 1 volume of IP dilution buffer (10 mM Tris pH 8.0, 1 mM EDTA, 200 mM NaCl, 1LmM EGTA. 0.2% Na-DOC, 1% Na-laurylsarcosine and 2% Triton X-100). 50Lµl of sheared chromatin was reserved as input and ChIP was performed overnight at 4 °C with rotation with 7.5 µg of H3K36me3 antibody (ab9050). 100ul Protein A dynabeads per sample were washed 3 times with 1ml chilled block buffer (0.5% BSA in PBS) and then added to the chromatin after overnight incubation with antibody and rotated for 4hr at 4 °C. Samples were washed 5 times in 1ml pre-chilled wash buffer (50mM HEPES pH 7.5, 500mM LiCl, 1mM EDTA, 1% IPEGAL-CA630, 0.7% Na-DOC and then 1ml pre-chilled TE + 50mM NaCl. Samples were eluted in elution buffer (50mM Tris pH 8.0, 10mM EDTA, 1% SDS) at 65°C. NaCl was added to final concentration of 455mM. Samples were incubated with 0.2mg/ml proteinase K for 1hr at 55°C and then decrosslinked overnight at 65°C. Samples were treated with 0.2mg/ml RNAase for 2hr at 37°C and then purified with the Zymo ChIP DNA clean and concentrator kit (D2505). Libraries were prepared using the NEBNext Ultra II DNA library prep kit (E7645) and sequenced by NovaSeq PE150. The sequence data were trimmed by Trimmomatic^22^ (v0.36) to remove adapter and then mapped to the hg38 assembly of the human genome using Bowtie2^19,20^ with the following settings: --local --very-sensitive --phred33 -X 1000. Reads with MAPQ values less than 10 were filtered using SAMtools (v1.8). Duplicate reads were removed using picard-tools. CHIP-seq signal was convereted to the bigwig format for visualization using deepTools bamCoverage^18^ (v3.3.1) with the following parameters: --binSize 10 --normalizeUsing CPM --effectiveGenomeSize 3209286105 --exactScaling.

### Immunofluorescence and DNA FISH staining

Coverslips were coated with 100 µg/ml poly-L-lysine overnight or 10 µg/ml laminin for 1hr at 37 °C before seeding cells. Asynchronized cells were seeded onto slides and subject to different treatment. Where indicated, edu was added to each well at 10 µg/ml 30 min before collecting samples. Immunofluorescence and dual-immunofluorescence DNA FISH staining were performed as described before. Briefly, slides were fixed with ice-cold 4% PFA for 15min, followed by permeabilization with 0.5% Triton X-100 in PBS for 15min at room temperature. Samples were blocked with 3% BSA in PBS for 1 hr at room temperature before incubation with primary antibody diluted in blocking buffer at 4°C overnight. After washing with PBS for a total of 3 times with 5 min each, slides were incubated with secondary antibody diluted in blocking buffer at room temperature for 1 h. Samples were fixed with ice-cold 4% PFA for 20 min after washing with PBS. If combined with DNA FISH staining, fixed samples were further permeabilized with ice-cold 0.7% Triton X-100 / 0.1M HCl (diluted in PBS) for 10 mins on ice. DNA was denatured by 1.5 M HCl for 30 mins at room temperature, followed by dehydration in ascending ethanol concentration. Diluted FISH probes (Empire Genomics) were pre-denatured at 75°C for 3min and added onto air dried slides. After incubation at 37 °C overnight, slides were washed with 2XSSC to get rid of non-specific binding, followed by DAPI staining. Where indicated, edu staining was performed with the Click-iT Plua EdU Alexa Fluor 647 Imaging kit (Invitrogen, cat:C10640).

### Replication Combing Assay (RCA)

Replication Fork Speed in extrachromosomal DNA (ecDNA) was evaluated using the molecular combing assay. COLO320DM and COLO320HSR cells were seeded into plates and let them grow into log phase, nascent DNA synthesize was pulse labelled with thymidine analogs: CldU and IdU sequentially for equal amount of time. Following pulse labeling, cells were harvested and embedded into agarose plugs using the Genomic Vision FiberPrep® kit (Genomic Vision, Bagneux, France). DNA extraction, combing and immunostaining was performed according to the EasyComb service procedures (Genomic Vision, Bagneux, France). Coverslips were scanned with FiberVision® scanner and images were analyzed using FiberStudio® software (Genomic Vision, Bagneux, France). Fork Speed was calculated using replication signals with contiguous CldU-IdU tracks. Only intact signals, flanked by counterstaining of the DNA fiber, were selected for analysis.

### Locus-Replication Combing Assay (Locus-RCA)

DNA replication activity at the MYC loci was assessed using molecular combing assay. COLO320DM and COLO320HSR cells were seeded into plates and let them grow into log phase, nascent DNA synthesize was pulse labelled with thymidine analogs: CldU and IdU for equal amount of time. Following pulse labeling, cells were harvested and embedded into agarose plugs using the Genomic Vision FiberPrep® kit (Genomic Vision, Bagneux, France). DNA extraction and combing was performed according to the EasyComb service procedures (Genomic Vision, Bagneux, France). DNA labeled FiberProbes® (Genomic Vision, Bagneux, France) targeting MYC loci were produced and hybridized to combed DNA. Correspondence between theoretical and experimental probe coverage patterns was validated by measuring hybridized probe length in control samples. After immunostaining of replication signals and DNA probes, coverslips were scanned with FiberVision® scanner. Image analysis and measurements were performed using FiberStudio® software (Genomic Vision, Bagneux, France). Fork Speed was calculated using replication signals with contiguous CldU-IdU tracks.

### Comet-FISH

Alkaline comet-FISH assays were performed according to, with minor modifications^41,42^. Cells were harvested by trypsinization, washed with PBS, and placed on ice. Cells were diluted in 37°C LMP agarose (IBI Scientific) in PBS to a final concentration of 0.7% and spread on pre-coated glass slides with a cover slip. Overnight lysis was performed at 4°C in alkaline lysis solution (2.5 M NaCl, 100 mM EDTA, 10 mM Tris pH 10, 1% TritonX-100, 10% DMSO) protected from light. The following day, slides were equilibrated for 30 min in alkaline electrophoresis buffer (200 mM NaOH, 1 mM EDTA, pH > 13) in a Coplin jar and subsequently electrophoresed at 25 V for 30 min. Slides were then neutralized with Tris, dehydrated in 70% ethanol, and dried at room temperature.

To detect ecDNA through FISH, Cy5-labeled probes were generated from RP11-440N18 BAC DNA sonicated to 150 bp and labeled using a DNA labeling kit (Mirus Bio). Slides were denatured with 0.5 M NaOH for 30 min at room temperature, dehydrated in an ethanol series (70%, 85%, 95%), and allowed to dry at room temperature. The hybridization mixture containing probe DNA (200 ng per slide) and Cot-1 DNA (8 μg per slide) was denatured separately at 75°C 10 min and pre-annealed at 37°C for 1 hr. Probe was added to the slides and spread with a glass cover slip and incubated at 37°C overnight in a humidified chamber. The following day, slides were washed four times in 2X SSC, 50% formamide at 42°C and subsequently washed twice in 2X SSC at 42°C. Slides were dipped briefly in 70% ethanol and air dried. Slides were mounted with Everbrite (Biotium) containing SYBR Gold (Invitrogen) diluted 1:10,000 and sealed with nail polish. Images were collected on a Nikon Eclipse TE2000-E using a 60X oil objective.

### Cell viability assay

Cell viability assay was performed using CellTilter-Glo (Promega, G8461) as previously described^43^. Briefly, cells were seeded into 384-well plate one day before adding inhibitors. Equal volumes of vehicles or drugs diluted at indicated concentration were added into each well the next day, and the cells were incubated for 3 days. On the third day, after equilibrating plate and CellTiter-Glo reagent at room temperature for 30 min, reagent was added into each well and incubated for 15 min at room temperature. Luminescence was measured using a synergy 2 microplate reader. Four biological replicates were performed for each condition. Data analysis was performed with GraphPad Prism.

### TUNEL

TUNEL assay (Invitrogen, C10617) was performed to detect DNA fragmentation during apoptosis. COLO320DM, COLO320HSR and SNU16 cells were treated with 1µM of CHIR-124 for indicated times. All cells including floating cells were collected and spun down onto slides using a cytospin (Thermo Scientific Cytospin 4 Centrifuge). Slides were fixed with 4% PFA and permeabilized with 0.25% TritonX-100, followed by labeling of free double strand end with EdUTP by reaction catalyzed by TdT enzyme in a humidified chamber at 37°C for 60 min. Incorporated EdUTP was detected through Click-iT™ Plus TUNEL reaction according to manufacturer’s manual at 37°C for 30 min. Slides were counter stained with DAPI and mounted with prolong-diamond antifade.

### Annexin V staining

Cell apoptosis was detected through flow cytometry using a FITC Annexin V Apoptosis Detection kit (BD bioscience, 556547). Cells were treated with 1µM of CHIR-124 for the indicated time, and all the cells including floating cells were collected. After washing with PBS twice and cell number counting, cells were resuspended in 1X binding buffer provided by the kit in a concentration of 1 × 10e6 cells/ml. 100 ml of the cell suspension was transferred to a FACS tube and stained with FITC Annexin V and PI. After incubation at RT for 15 min, all the samples were analyzed with BD LSRII follow cytometry (BD Biosciences) within 1 hr.

### Microscope and Image analysis

Images were taken by conventional fluorescence microscopy or confocal microscopy. Conventional fluorescence microscopy was performed on a Leica DMi8 widefield microscope using a 63× oil objective. Confocal microscopy was performed on ZEISS LSM 880 inverted confocal microscope (Stanford CSIF Facility). A z stacks were taken for each field of view and a best-in-focus stack was identified for downstream image analysis except for Fig 2d and Fig 3a, where a max projection was performed by ImageJ.

Image analysis and quantification were performed using the open-source software CellProfiler. For foci number analysis, DAPI staining, IF staining and DNA FISH channel were analyzed through automatic thresholding and segmentation to call nuclei, pRPA2S33/γH2AX foci and DNA FISH foci respectively. Co-localization was performed using an object-based co- localization method. For fluorescence intensity measurement, nuclei were called based on DAPI channel through automatic thresholding and segmentation, mean fluorescence intensity was retrieved from measuring mean fluorescence intensity within each nucleus.

### Replication stress score computation

The replication stress signature score of each sample from TCGA was retrieved from literature from Llobet et al^21^, which were transformed linearly between zero and one by subtracting the minimum score and dividing by the maximum score. TCGA sample ecDNA status classification was performed as stated in previous publication^20^. Briefly, 1921 TCGA samples were grouped into five sub-types by Amplicon Classifier: ecDNA, BFB, complex non-cyclic, linear, and no-amplification. Samples with a BFB or complex non-cyclic status were removed from the analysis due to the challenges of detecting ecDNA from short read data. Samples with linear amplification and no-amplification were classified as ecDNA-. After removing metastasis sample and ecDNA- samples without matching ecDNA+ samples in the same tissue origin, a total of 234 ecDNA+ and 636 ecDNA- samples were included in the analysis.

### CRISPR experiment

sgRNA template oligos targeting gene encoding CHK1 was synthesized (Integrated DNA technologies IDT) and was ligated into a CRISPR expression vector with RFP (Cellecta- pRSG16-U6-sg-HTS6C-UbiC-TagRFP-2A-Puro). Non-targeting GFP (sgNT-GFP) plasmid was purchased.

ecDNA+ and ecDNA- Hela cells were transduced with sgCHK1-RFP or sgNT-GFP virus, and puromycin (Sigma) was added at 2.5 µg/ml for selection for 48 hours. After 48 hours of puromycin selection (Day 0), equal number of cells expressing either sgCHK1-RFP and sgNT-GFP were mixed to obtain the RFP/GFP population ratio. In the following days, flow cytometry analysis was performed to determine the sgCHK1-RFP/sgNT-GFP ratio. The mixed cell population cultures were maintained at sub-confluency. sgRNA sequence targeting CHK1 were as follows:

#17: CCTGACAGCTGTCACTGGGT

#18: GCTGTCAGGAGTATTCTGAC

### Western blotting

Samples were lysed in Radioimmunoprecipitation (RIPA) assay buffer (Boston BioProducts BP- 115) supplemented with protease/phosphatase inhibitors (Fisher Scientific 78444). Protein concentration was quantified with Bicinchoninic acid assay (BCA) assay (Fisher Scientific 23225) and samples were prepared in 4X sample buffer (Bio-Rad 1610747). Samples were loaded and run on 4-12% Bis-Tris Gradient Gel (Fisher Scientific WG1403BOX) and transferred onto nitrocellulose membrane (Bio-Rad 1704271). The membrane was blocked with 5% BSA in Tris-buffered saline with tween (TBST; Fisher Scientific 28360) for an hour and then primary antibody was added and incubated overnight at 4°C. Following primary antibody incubation, the membrane was washed with TBST and incubated with secondary antibody for 1h. The membrane was then incubated with Enhanced chemiluminescence reagent (ECL; Fisher Scientific 32106) and image acquisition was performed on ProteinSimple FluorChemE.

### Detection of phosphorylated CHK1 Ser345 using the AlphaLisa Sure Fire assay

Compound activity in cells was measured using an AlphaLISA® SureFire® Ultra™ p-CHK1 (Ser345) assay (Perkin Elmer, catalog no. ALSU-PCHK1-A10K). HT29 cells were cultured in McCoy 5A medium with 10% FBS and 1% penicillin-streptomycin and seeded to 96-well plates (Corning, catalog no. 3599). Compounds were serially diluted in DMSO over a 10-point dose range with 3-fold dilution and to each well containing cells was added compound solution. Plates were centrifuged at 1000 rpm for 30 seconds. Plates were incubated at 37 °C for 16 h. Supernatant was removed by flicking the plate against a paper towel. Wells were washed once with PBS solution. To each well was added freshly prepared lysis buffer and plates were agitated on a plate shaker at 400 rpm for 30 min. The 96-well cell plates were centrifuged at 1500 rpm for 1 minute. From each well was transferred 10 µL of the lysates to a 384-well Optiplate™ (Perkin Elmer, catalog no. 6007290). To each well was added Acceptor Mix (5 µL) and the plates were sealed and wrapped in foil. Plates were agitated on a plate shaker for 2 minutes, then incubated at room temperature for 1 h. To each well was added Donor Mix (5 µL) and the plates were sealed and wrapped in foil. Plates were agitated on a plate shaker for 2 minutes, then incubated at room temperature for 1 h. AlphaLisa signal was read on an EnVision multimode plate reader (Perkin Elmer). Data were fitted to dose-response curves using XLfit (IDBS, Surrey, UK) or GraphPad Prism (GraphPad Software, La Jolla, CA, US) to calculate IC50 values for each compound tested.

### Kinase HTRF biochemical assay

Chk1 enzyme activity was measured using an HTRF KinEASE assay (Cisbio, catalog no. 62ST1PEC). Full-length human CHK1 protein (GenBank accession number NP_001265.1) was obtained from Carna Biosciences, Inc. (Kobe, Japan, catalog no. 02-117). The enzyme reaction was carried out in assay buffer containing (final concentrations): CHK1 enzyme (0.012 ng/µL), MgCl2 (5 mM) and DTT (1 mM). To determine compound dose response, DMSO stock solutions were serially diluted in a 10-point concentration series in duplicate. Compound solution (50 nL) was added to 384-well assay plates (Greiner, catalog no. 784075). To each well containing compound solution was added assay buffer solution (5 µL). Plates were centrifuged at 1000 rpm for 1 minute, then incubated at room temperature for 10 minutes. The reaction was started by addition of substrate buffer (5 µL/well) containing (final concentrations): STK substrate 1-biotin (120 nM) and ATP (1 mM). Assay plates were centrifuged at 1000 rpm for 1 minute, then incubated at room temperature for 60 minutes. The reaction was stopped by addition of detection buffer (Cisbio, 10 µL) containing (final concentrations): STK antibody- cryptate (0.25 nM) and streptavidin-XL665 (7.5 nM). Plates were centrifuged at 1000 rpm for 1 minute, then incubated at 25 °C for 2 hours. HTRF signal was read on an EnVision multimode plate reader (CisBio) in HTRF mode. Data were fit to dose-response curves using XLfit (IDBS, Surrey, UK) or Prism (GraphPad Software, La Jolla, CA, US) to calculate IC50 values for each compound tested.

### Phospho-RPA32 S8 Immunofluorescence High Content Imaging

Optical-bottom 96-well plates (Thermo Scientific, #165305) were coated with 50 µL of 1:1 Poly-L-Lysine (R&D Systems, #3438-100-01) and Poly-D-Lysine (R&D Systems, #3439-100-01) for 3 h at RT. The wells were washed once with 100 µL of phosphate buffered saline (PBS) (Gibco, #10010-023) and all liquid was removed from the wells and allowed to dry fully at RT. COLO320 ecDNA+ cells were seeded at 15,000 cells per well in 100 µL of RPMI media (Thermo Fisher, #22400089) supplemented with 10% FBS (Omega Scientific, #FB-01). Cells were left to attach in a 37 °C incubator with 5% CO2 overnight. The following day, cells were treated with BBI-825 for 16 h. Following treatment, all culture media was removed, and cells were fixed with 4% paraformaldehyde (PFA) (Boston BioProducts, #BM-155) for 15 min at RT. After fixation, the 4% PFA was removed, and wells were washed twice with 100 µL of PBS. The cells were then permeabilized with 100µL of 0.5% Triton X-100 (Sigma-Aldrich, #T8787) in PBS for 15 min at RT. After permeabilization, wells were washed twice with 100 µL of PBS and then blocked with 5% goat serum (Abcam, #ab7481) and 1 mg/mL of BSA (GeminiBio, #700- 100P) for 1 h at RT. The primary antibody (phospho-RPA32 (S8); Cell Signaling, #54762) was diluted at 1:200 in blocking buffer and 50 µL was added to all wells and incubated at 4 °C overnight. Plates were then washed three times with 100 µL of PBS and then incubated with 1:1000 dilution of secondary antibody (Goat anti-rabbit IgG Alexa Fluor Plus 594; Thermo Fisher, #A32740s) and 1:1000 dilution of Hoechst 33342 (Biotium, #40046) in blocking buffer for 1 h at RT. Plates were then washed three times with 100 µL of PBS, 100 µL of PBS left in the wells following the final wash. The plate was imaged using a CellInsight CX7 LZR Pro high content imager (ThermoFisher Scientific, Waltham, MA) and data analyzed using the Spot Detector BioApplication module on the HCS Studio Cell Analysis software (ThermoFisher Scientific, Waltham, MA). Puncta were detected using a pixel thresholding method within a nucleus and cells that contained 3 or more puncta of phosphorylated RPA32 Ser8 staining were considered as a positive signal.

### Xenograft

Animal experiments were performed in accordance with protocols approved by the CRADL Institutional Animal Care and Use Committee (Protocol #EB17-010-066). SNU-16 gastric cancer cell line was purchased from ATCC (ATCC #CRL5974) and maintained in RPMI growth medium (Gibco #22400-089) supplemented with 10% fetal bovine serum (FBS) (Omega Scientific #FB-02). To establish tumors, 1L×L10e6 SNU-16 cells in 200 µL of a 1:1 mixture of PBS and Matrigel (Corning #354234) by subcutaneous injection into the right flank of 9-week-old female SCID beige mice (strain code 186; Envigo, Livermore, CA). Tumor measurements were taken two-three times per week, body weights were taken daily. Tumor volume measurements were obtained using digital calipers and tumor volumes (TV; mm3) were determined using the formula TV = (L × W2)/2 where L is the length/largest tumor diameter and W is the width/shortest tumor diameter. Animals (eight mice per group) were randomly assigned to treatment with vehicle, infigratinib (15 mg kg-1 PO QD), BBI-2779 (30 mg kg-1 PO Q2D), or the combination of BBI-2779 and infigratinib once average tumor volume was 286 (+/- 10) / mean (+/-SEM) mm3. Infigratinib was formulated in a 1:1 mixture of sodium acetate buffer, pH 4.6 and polyethylene glycol (PEG) 300. BBI-2779 was formulated in 0.5% methylcelluose (Sigma Aldrich, #M0512) and 0.2% Tween 80 (AG Scientific, #T-2835) in HyPure Molecular Biology Grade Water (HyClone, #SH30538.02). Dose holidays were provided to individual animals that demonstrated > -10% body weight change from baseline and Nutra-Gel was provided to entire treatment group. Animals were sacrificed 6-hours, 24-hours, or 36 hours after the last dose and tumors were collected for western blot or copy number analysis.

### Copy Number Analysis from xenograft samples

For copy number analysis, tumors were cut into 10-20 mg pieces and flash frozen in liquid nitrogen. DNA was extracted using the QIAcube DNA Extraction Kit (Qiagen, #51331). Briefly, a mixture of Buffer ATL and Proteinase K was added to the frozen tumor pieces, and they were set out to equilibrate to RT. Tumors were then vortexed for 30 sec and placed into an incubator at 56 °C to digest overnight. The next morning an additional 150 ul of Buffer ATL was added and samples vortexed an additional 30 sec to reduce viscosity of the samples before transfer to the S block. Qiagen protocol for the 96 QIAcube HT was followed for the remainder of the DNA isolation. Purified DNA was quantified for the presence of dsDNA on the QIAxpert (Qiagen #9002340). The DNA was diluted to 5 ng/µL (5x working stock) in RNase/DNase free water (Thermo Fisher Scientific #10977015) and 2 µl was loaded into a 384-well plate. Master mix recipe (Master Mix (2x), 5.5 µl; CNA (Target Gene) 20x, 0.55µl; CNR (TERT) 20x, 0.55 µl; Nuclease-free water, 2.2 µl) was made containing TaqPath Pro MasterMix 2x (Thermo Fisher Scientific, #A30866) human Female genomic DNA (Promega #G1521), as a reference, internal controls (human Tert) and FGFR2 or MYC target gene probe (Thermo Fisher Scientific #4400292). Reactions were run on the QuantStudio 6 or 7 (ThermoFisher Scientific, Waltham, MA) using the qPCR reaction settings as follows: Denature/Enzyme activation: 95**°**C, 10 min; 40 cycles of Denature 95 **°**C 15 sec, Anneal/Extend 60**°**C 60 sec.

### Synthesis of BBI-2279

All chemicals were purchased from commercial suppliers and used as received unless otherwise indicated. Proton nuclear magnetic resonance (1H NMR) spectra were recorded on Bruker AVANCE 400 MHz spectrometers. Chemical shifts are expressed in δ ppm and are calibrated to the residual solvent peak: proton (CDCl_3_, 7.26 ppm). Coupling constants (J), when given, are reported in hertz. Multiplicities are reported using the following abbreviations: s = singlet, d = doublet, dd = doublet of doublets, t = triplet, q = quartet, m = multiplet (range of multiplet is given), br = broad signal, and dt = doublet of triplets. Carbon nuclear magnetic resonance (13C NMR) spectra were recorded using a Bruker AVANCE HD spectrometer at 100 MHz. Chemical shifts are reported in parts per million (ppm) and are calibrated to the solvent peak: carbon (CDCl3, 77.23 ppm).

All final compounds were purified by reverse phase HPLC or SFC. The purity for test compounds was determined by HPLC on a SHIMADZU LC-2010A HT instrument. HPLC conditions were as follows: XBRIDGE C18 3.5um 2.1*50mm, H2O(0.05%TFA)- ACN(0.05%TFA), ACN from 0 to 60% over 7 minutes, 7-8min, ACN from 60% to 100%, flow rate 0.8 mL/min, UV detection (λ =214, 254 nm). The mass spectra were obtained using LCMS on a LCMS-Agilent 6125 instrument using electrospray ionization (ESI). LCMS conditions were as follows: Column: Waters Cortecs C18+, 2.7um 30 mm; Mobile phase : ACN (0.05% FA)-Water (0.05% FA); Gradient: 5% ACN to 95% ACN in 1.0 min, hold 1.0 min, total 2.5 min; Flow 1.8 mL / min; UV detection (λ = 214, 254 nm). Column Temp:45 degree. The SFC purity for test compounds was determined with a SFC Thar prep 80

#### Step 1: 2-bromocyclobutan-1-one (**2**)

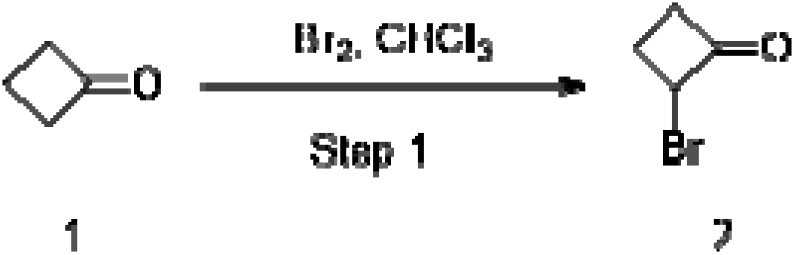

To a solution of cyclobutanone (10 g, 0.142 mol) in CHCl_3_ (100 mL) was was added Br_2_ (22.82 g, 0.170 mmol) at 0 °C. The reaction mixture was stirred at 25 °C for 12 hours. The residue was quenched by saturated aqueous sodium thiosulfate (100 mL) and extracted with ethyl acetate (100 mL × 3). The organic layers were combined and dried over Na_2_SO_4_ and concentrated to give 2-bromocyclobutan-1-one (19.5 g, 91.9% yield) as a colorless oil, which was used in the next step without further purification.

#### Step 2: 2-(2-bromo-3-methoxyphenoxy)cyclobutan-1-one (3)

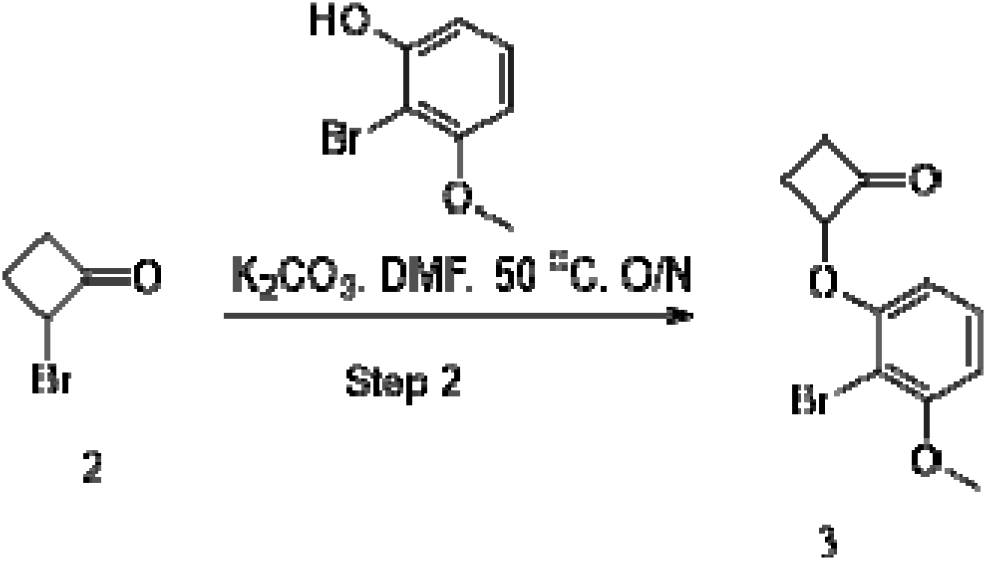

A solution of 2-bromo-3-methoxyphenol (3 g, 0.016 mol), 2-bromocyclobutan-1-one (7.068 g, 0.080 mol) and K_2_CO_3_ (5.447 g, 0.040mol) in DMF(30 mL) was stirred at 50 °C for 12 hours. The mixture was filtered and concentrated under reduced pressure. After concentration, the residue was purified via flash column chromatography, eluted with petroleum ether/ethyl acetate (from 0% to 40%) to give 2-(2-bromo-3-methoxyphenoxy)cyclobutan-1-one (1.65 g, 42.3 % yield) as a white solid. MS (ESI): mass calcd. for C_11_H_11_BrO_3_ 269.99, 271.99 m/z found 270.9, 272.9 [M+H]^+^. LCMS (method 1, 2.5 min formic acid): Rt = 1.303 min

#### Step 3: N-((1R,2R)-2-(2-bromo-3-methoxyphenoxy)cyclobutyl)-2-methylpropane-2-sulfinamide (4)

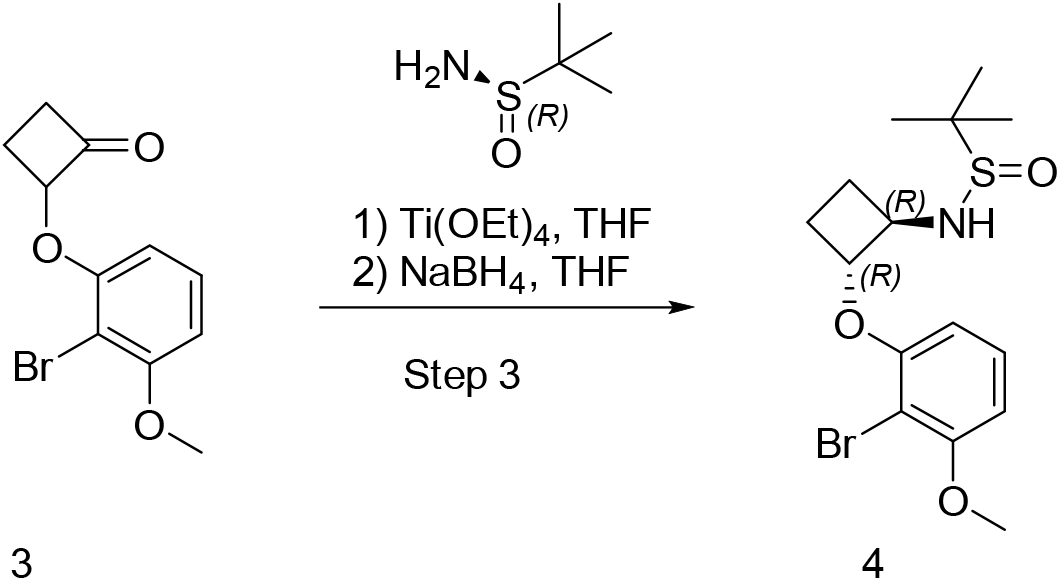

To a solution of 2-(2-bromo-3-fluorophenoxy)cyclobutan-1-one (200 mg, 0.7720 mmol), (S)-2-methylpropane-2-sulfinamide (102.92 mg, 0.8492 mmol) and Ti(OEt)_4_ (329.13 mg, 1.158 mmol) in THF was stirred under nitrogen for 2 hours then NaBH_4_ (58.41 mg, 1.544 mmol) wa added at 0 °C. The reaction mixture was stirred at 50 °C for 2 hours. The reaction was quenched by MeOH (5 mL) and H_2_O (50 mL), then extracted with ethyl acetate (50 mL × 3). The organic layers were combined and dried over Na_2_SO_4,_ and concentrated under reduced pressure at 30 °C. The residue was purified by Prep-HPLC (Daisogel-C18-10-100, 30 × 250 mm, 5 um, mobile phase: ACN--H2O (0.1%FA), gradient: 5 ∼ 95) to afford 1: N-((1R,2R)-2-(2-bromo-3- methoxyphenoxy)cyclobutyl)-2-methylpropane-2-sulfinamide (30 mg, 12.1 % yield) as a white solid. MS (ESI): mass calcd. for C_15_H_22_BrNO_3_S 375.05, 377.05, m/z found 376.0, 378.0[M+H]^+^. LCMS (method 1, 2.5 min formic acid): Rt = 1.320 min.

#### Step 4: tert-butyl3-((tert-butoxycarbonyl)(5-cyanopyrazin-2-yl)amino)-5-(2-((1R,2R)-2- ((tert-butylsulfinyl)amino)cyclobutoxy)-6-methoxyphenyl)-1H-pyrazole-1-carboxylate (5)

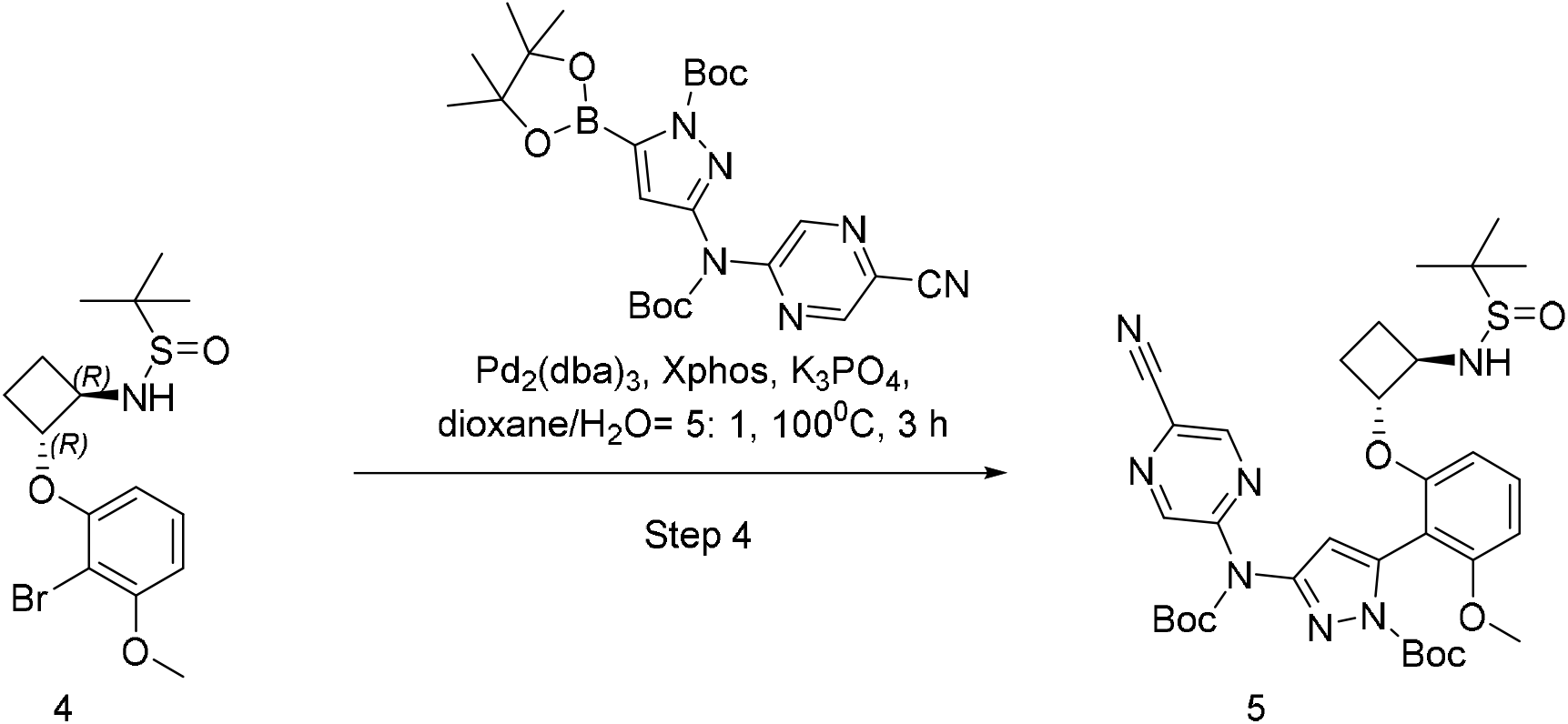

A solution of N-((1R,2R)-2-(2-bromo-3-methoxyphenoxy)cyclobutyl)-2-methylpropane-2- sulfinamide (200 mg, 0.5490 mmol), tert-butyl-3-((tert-butoxycarbonyl)(5-cyanopyrazin-2-yl)amino)-5-(4,4,5,5-tetramethyl-1,3,2-dioxaborolan-2-yl)-1H-pyrazole-1-carboxylate (338.87 mg, 0.6588 mmol) and K_3_PO_4_ (349.61 mg,1.6470 mmol) in dioxane : H_2_O = 5 : 1, (11 mL) was stirred under nitrogen at 100 °C. Then, X-phos (104.69 mg, 0.2196 mmol) and Pd_2_(dba)_3_ (100.55 mg, 0.1098 mmol) were added. The reaction mixture was stirred at 100 °C for 3 hours. The mixture was filtered and concentrated under reduced pressure. After concentration, the residue was purified via flash column chromatography, eluted with DCM/MeOH (from 0% to 10%) to give tert-butyl3-((tert-butoxycarbonyl)(5-cyanopyrazin-2-yl)amino)-5-(2-((1R,2R)-2-((tert- butylsulfinyl)amino)cyclobutoxy)-6-methoxyphenyl)-1H-pyrazole-1-carboxylate (150 mg, 40.7 % yield) as a orange solid. MS (ESI): mass calcd. for C_33_H_43_N_7_O_7_S 681.29, m/z found 682.2 [M+1]^+^. LCMS (method 1, 2.5 min formic acid): Rt = 1.402min

#### Step 5: 5-((5-(2-((1R,2R)-2-aminocyclobutoxy)-6-fluorophenyl)-1H-pyrazol-3- yl)amino)pyrazine-2-carbonitrile (BBI-2779)

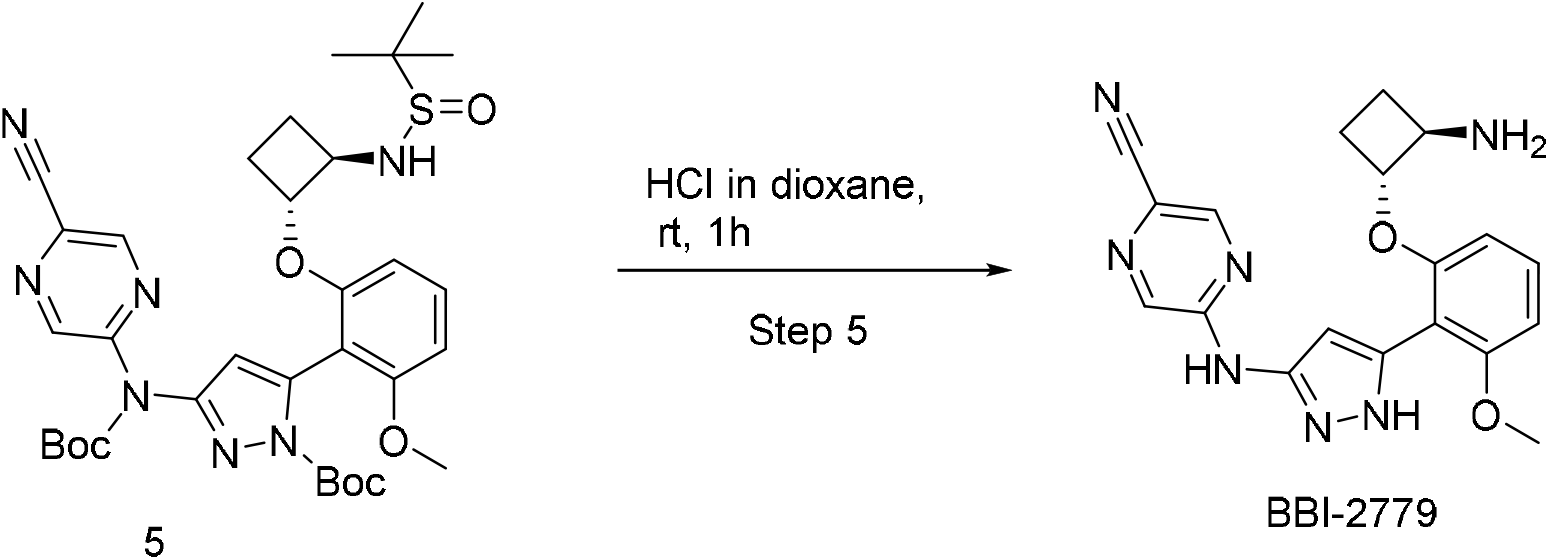

To a solution of tert-butyl3-((tert-butoxycarbonyl)(5-cyanopyrazin-2-yl)amino)-5-(2- ((1R,2R)-2-((tert-butylsulfinyl)amino)cyclobutoxy)-6-fluorophenyl)-1H-pyrazole-1-carboxylate (200 mg, 0.3505 mmol) in HCl in dioxane (4M, 10 mL) was stirred at 25 °C for 1 hour. It was the concentrated under reduced pressure. The residue was purified by Prep-HPLC (Daisogel- C18-10-100, 30 × 250 mm, 5 um, mobile phase: ACN--H2O (0.1%FA), gradient: 5 ∼ 95) to give 5-((5-(2-((1R,2R)-2-aminocyclobutoxy)-6-methoxyphenyl)-1H-pyrazol-3-yl)amino)pyrazine-2- carbonitrile (25 mg, 19.5% yield) as a white solid. MS (ESI): mass calcd. for C_19_H_19_N_7_O_2_ 377.16, m/z found 378.0 [M+H]^+^. LCMS (method 1, 2.5 min formic acid): Rt = 1.005 min ^1^H NMR (400 MHz, DMSO-*d*_6_) δ ppm 10.72 (s, 1H), 8.65 (d, *J* =0.8 Hz, 1H), 8.57 (s, 1H), 8.33 (s, 1H), 7.30 (1 H, t, *J* = 8.4 Hz, 1H), 6.93 (s, 1H), 6.76 (t, *J =* 8.8 Hz, 1H), 4.66-4.46 (m, 1H), 3.82 (s, 3H), 3.59-3.56 (m, 1H), 2.35-2.30 (m, 1H), 2.12-2.05 (m, 1 H), 1.62-1.49 (m, 2H).

### Quantifications and statistical analysis

All statistical methods and sample size have been stated in figure legend or method section. No statistical methods were used to predetermine the sample size. The default test type was a two-sided statistic test, unless indicated in the text. The investigators were not blinded to allocation during experiments and outcome assessment.

